# Generation of human alveolar epithelial type I cells from pluripotent stem cells

**DOI:** 10.1101/2023.01.19.524655

**Authors:** Claire L Burgess, Jessie Huang, Pushpinder Bawa, Konstantinos-Dionysios Alysandratos, Kasey Minakin, Michael P Morley, Apoorva Babu, Carlos Villacorta-Martin, Anne Hinds, Bibek R Thapa, Feiya Wang, Adeline M Matschulat, Edward E Morrisey, Xaralabos Varelas, Darrell N Kotton

**Affiliations:** Center for Regenerative Medicine of Boston University and Boston Medical Center, Boston, MA 02118, USA; The Pulmonary Center and Department of Medicine, Boston University Chobanian & Avedisian School of Medicine, Boston, MA 02118, USA; Penn-CHOP Lung Biology Institute, Perelman School of Medicine, University of Pennsylvania, Philadelphia, PA, USA; Department of Biochemistry, Boston University Chobanian & Avedisian School of Medicine, Boston, MA, 02118, USA

## Abstract

In the distal lung, alveolar epithelial type I cells (AT1s) comprise the vast majority of alveolar surface area and are uniquely flattened to allow the diffusion of oxygen into the capillaries. This structure along with a quiescent, terminally differentiated phenotype has made AT1s particularly challenging to isolate or maintain in cell culture. As a result, there is a lack of established models for the study of human AT1 biology, and in contrast to alveolar epithelial type II cells (AT2s), little is known about the mechanisms regulating their differentiation. Here we engineer a human in vitro AT1 model system through the directed differentiation of induced pluripotent stem cells (iPSC). We first define the global transcriptomes of primary adult human AT1s, suggesting gene-set benchmarks and pathways, such as Hippo-LATS-YAP/TAZ signaling, that are enriched in these cells. Next, we generate iPSC-derived AT2s (iAT2s) and find that activating nuclear YAP signaling is sufficient to promote a broad transcriptomic shift from AT2 to AT1 gene programs. The resulting cells express a molecular, morphologic, and functional phenotype reminiscent of human AT1 cells, including the capacity to form a flat epithelial barrier which produces characteristic extracellular matrix molecules and secreted ligands. Our results indicate a role for Hippo-LATS-YAP signaling in the differentiation of human AT1s and demonstrate the generation of viable AT1-like cells from iAT2s, providing an in vitro model of human alveolar epithelial differentiation and a potential source of human AT1s that until now have been challenging to viably obtain from patients.

## Introduction

The alveolar epithelium of the lung is vital for gas exchange and consists of two cell types, referred to as alveolar epithelial type I (AT1) and type II (AT2) cells. AT2s are cuboidal and produce surfactant, while AT1s are uniquely flattened to allow for the diffusion of oxygen into the capillaries. AT1s make up 40% of the alveolar epithelial cell population but cover over 95% of the alveolar surface area.^1^ Despite their critical role comprising the vast majority of the lung’s alveolar surface area and recent revelations of the role of AT1 differentiation in fibrotic lung disease,^2–4^ relatively little is known about the cell biology, origins, and fates of human AT1s.

The expansive and fragile structure of AT1s has made them particularly challenging to study. A single AT1 can cover the surface of multiple adjacent alveoli, making it difficult to identify individual AT1s with a single 2D cross-section.^1, 5^ Likewise, AT1s are easily shredded by methods such as cell sorting, leading to difficulties in isolating fresh primary AT1s. Their putative terminally differentiated nature has limited efforts to maintain these cells in vitro with only a few prior reports suggesting successful expansion in culture as proliferating cells.^6, 7^

AT1s are generally assumed to be replenished by adjacent cuboidal AT2s, based on a broad literature beginning in the 1970’s that employed rodent models of lung injury to characterize the transition of AT2 progenitor cells into AT1 progeny in vivo^8, 9^ or after culturing in vitro.^3, 4, 10, 11^ More recently, lineage tracing in mice has shown that AT1s can be derived from mature AT2s after injury and during normal homeostasis.^10, 12, 13^ The exact signaling mechanisms driving AT1 differentiation in adults are still uncertain, although recent studies in mice have implicated a wide variety of classical pathways such as Wnt, BMP, TGFb, FGF, and YAP/TAZ signaling.^14–21^ Additionally, mouse studies have shown that nuclear YAP/TAZ localization is necessary for the maintenance of the AT1 program, and that loss of YAP/TAZ in mature AT1 cells leads to reversion back into an AT2-like cell fate.^22, 23^

The origin and differentiation of AT1s in humans, however, has been less clear and more difficult to study. A variety of reports involving the in vitro 2D culture of isolated primary fetal or adult human AT2s have documented the rapid downregulation of AT2 marker transcripts and proteins, coincident with a flattened morphology, and upregulation of some purported AT1 markers of unclear specificity, leading to the currently accepted paradigm that AT1s are a “default state” of AT2s, if cultured in undefined (e.g. serum containing) conditions.^24–27^ This default state has been called into question by more recent 3D culture models which observed little if any evidence of bona fide AT1 cells emerging from either primary human AT2 cells co-cultured with^28^ or without fibroblasts or from human induced pluripotent stem cell (iPSC)-derived AT2s (iAT2s), even after prolonged time in most culture conditions tested to date,^29–31^ with three notable exceptions.^31–35^

Understanding the differentiation of human AT1s during homeostasis and after lung injury is particularly important since damage to AT1s in response to toxic inhalational exposures, radiation, or a variety of infections can lead to respiratory failure and severe diseases such as the Acute Respiratory Distress Syndrome (ARDS).^36^ Impaired differentiation of AT1s has also recently been implicated in fibrotic lung diseases.^3, 4, 37, 38^ Hence, understanding normal differentiation of human AT1s could provide insights for resolving the aberrant transitional state found in fibrotic lung tissue. The engineering of laboratory mice carrying fluorochrome reporters or lineage tracing cassettes under the regulatory control of AT1 gene promoters has been invaluable in identifying, tracing, and isolating mouse AT1s during development and disease;^6, 39, 40^ however, to date no comparable engineered human reporter has yet been generated to facilitate the study of human AT1s.

Here we report the in vitro generation of cells expressing the molecular and functional phenotypes of human AT1s via differentiation of iAT2s. We first profile human lung explant tissues at single cell resolution using RNA-sequencing to identify potential AT1-selective marker gene-sets and AT1-enriched signaling pathways. We engineer an iPSC line carrying a tdTomato reporter targeted to the endogenous AGER locus, which is specifically upregulated in primary AT1s according to our gene-set. The AGER^tdTomato^ reporter enables real time tracking and quantitation, of the resulting putative iPSC-derived AT1s (iAT1s), facilitating the isolation of a pure population of iAT1s. We find activated nuclear YAP expression is sufficient to drive the transcriptomic shift from iAT2 to iAT1 programs in a cell-autonomous manner in multiple iPSC lines. Further, we have developed a defined, serum-free differentiation medium containing a LATS inhibitor that recapitulates the above process, leading to robust and efficient differentiation of iAT2s into iAT1s.

## Results

### Single cell RNA sequencing defines the transcriptomic program of primary human alveolar epithelial type I cells

To understand the transcriptomic programs of primary human AT1s, we performed profiling by single cell RNA sequencing (scRNA-seq) of 5 distal human lung explant tissues (partial dataset published previously in Basil et al, *Nature* 2022)^41^. After visualization of all pooled analyses, we identified 19 distinct epithelial, mesenchymal, and immune cell clusters, including 1401 AT1 cells, and annotated each cluster according to canonical lineage markers detailed previously for this dataset (Fig. S1a).^41^ Selecting and reclustering all epithelial cells (UMAP; Fig. 1a), we visualized expression of previously reported putative AT1 markers (*PDPN, CAV1, AGER, AQP5,* and *HOPX*) and canonical AT2 marker genes (*SFTPC, LAMP3, ABCA3*) through UMAP overlays (Fig. 1b). We found some human AT1 markers (*AGER, CAV1, PDPN*, and *RTKN2*) shared similar qualitative clustered expression patterns to published mouse scRNA-seq atlases;^42, 43^ however, *HOPX* and *AQP5*, which have been well characterized as mouse AT1-specific markers,^6, 28^ lacked specificity as they were also expressed in other epithelial cells, such as AT2s for *HOPX* and airway cells for *AQP5*. (Fig. 1b-d, S1b).

**Figure 1:**
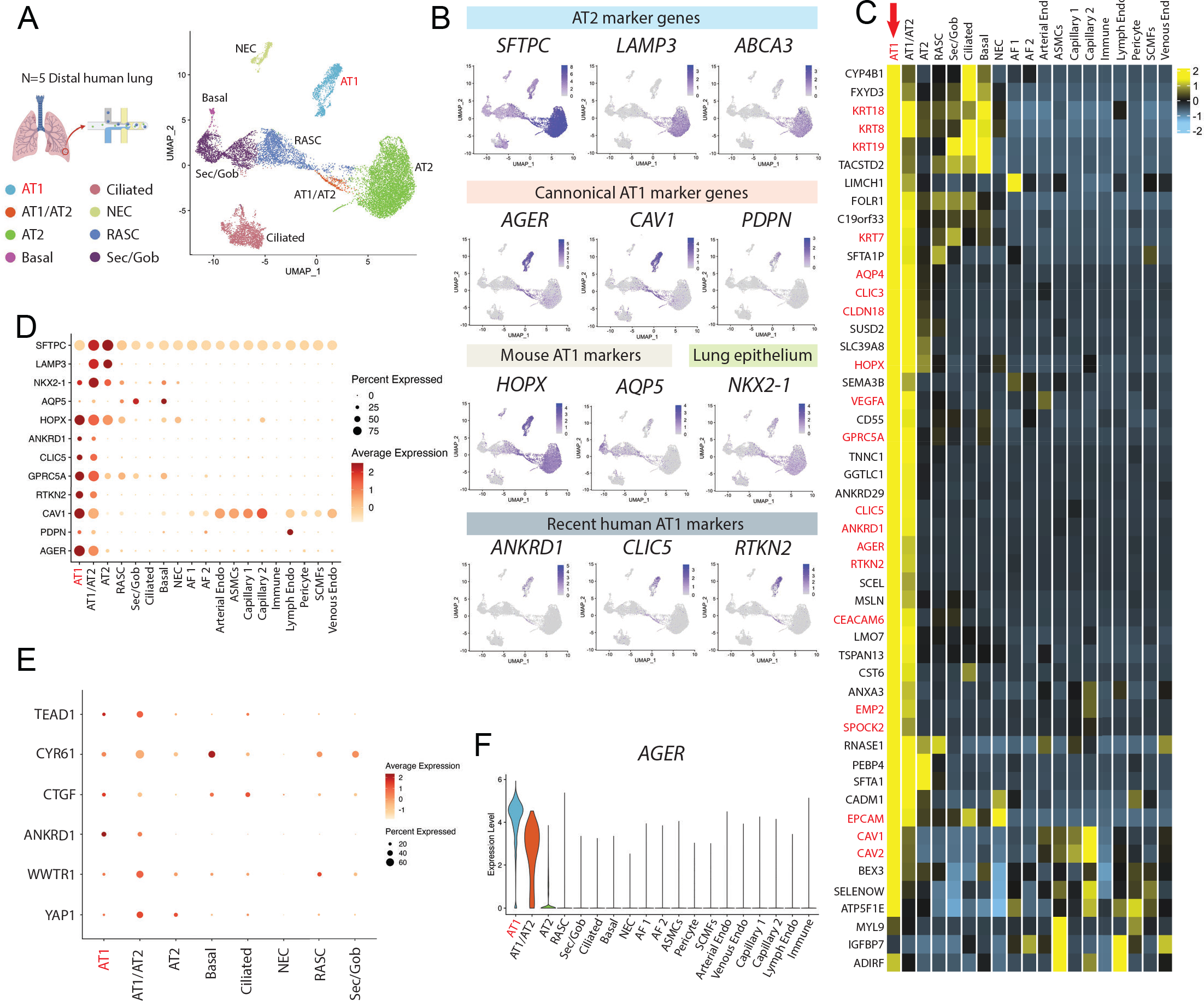
Transcriptomic profiling at single cell resolution of primary adult human AT1s. A) Uniform Manifold Approximation and Projection (UMAP) of an integrated analysis of 15769 primary distal lung epithelial cells (dataset previously published in Basil et al). N=5 including, 1401 AT1s and 7039 AT2s.^41^ B) Expression of AT2 specific marker genes, *SFTPC, LAMP3, ABCA3*; canonical AT1 marker genes *AGER, CAV1, PDPN*; mouse-specific AT1 marker genes *HOPX*, *AQP5*; lung epithelial marker *NKX2-1*; and more recent human AT1 marker genes *ANKRD1*, *CLIC5*, and *RTKN2*. C) Heatmap showing average expression for each cell type of top differentially upregulated human AT1 genes compared to all distal lung cells. D) Expression of selected AT1 and AT2 genes across all lung cell types. E) Expression of YAP (*YAP1*) and TAZ (*WWTR1*) and known downstream Hippo signaling markers across lung epithelial cell types. F) Violin plot showing *AGER* expression across indicated cell types. Also see supplemental figure 1.

To generate unbiased candidate human AT1 marker 50-gene sets we identified differentially expressed genes (DEGs) enriched in AT1s using three pair-wise comparisons: 1) AT1s vs all lung cells, 2) AT1s vs. lung epithelial cells, and 3) AT1s vs AT2s (Fig. 1c and Table S1). *AGER* was the top transcript significantly enriched in human AT1s in each of the 3 comparisons (ranked by logFC) with multiple caveolin (*CAV1* and *CAV2*) and chloride intracellular channel (*CLIC3* and *CLIC5*) gene family members present in the top upregulated 50 genes as well (Fig 1c and Table S1). Although *PDPN* was not in the top 50 upregulated genes, likely due to its expression in lymphatic endothelium and basal cells, we still selected it as an informative AT1 marker because *PDPN* was significantly upregulated in all comparisons, and *PDPN* has been extensively published as a canonical AT1 marker in other scRNA-seq atlases for both mice and humans.^42–46^ To confirm the utility of the above markers, we compared their expression levels, frequencies, and relative specificities across all human lung epithelia (Fig. 1d-f and S1b). Taken together, these analyses suggested *AGER, CAV1, CLIC5*, and *PDPN* as an informative 4-gene human AT1 marker set, in addition to the more extended 50-gene marker sets (Table S1), each able to reliably distinguish human AT1 cells.

We noted the canonical YAP/TAZ target gene, *ANKRD1*, was in the top 50 most enriched transcripts in AT1 cells in each of our comparisons (Table S1), a finding consistent with recent publications suggesting activated nuclear YAP/TAZ is important for the differentiation and maintenance of the mouse AT1 program.^22, 47–50^ Other downstream targets of YAP/TAZ, including *CTGF*, *CYR61*, and *TEAD1*, were also highly expressed in the AT1 population as well as in transitioning AT1/AT2 cells (Fig. 1e). However, *YAP1* and TAZ (*WWTR1*) mRNAs were expressed broadly and not specifically enriched in AT1s (Fig. 1e), an observation that is consistent with YAP/TAZ activity being primarily defined through post-translational regulation.^51^ Taken together these analyses: a) defined the transcriptomic programs of human AT1 cells, including multiple extended (50-gene) AT1 marker gene sets (Table S1), b) validated a quartet of canonical human AT1 transcript markers (*AGER, CAV1, CLIC5*, and *PDPN*), and c) suggested that YAP/TAZ activation distinguishes AT1 from AT2 cells in humans.

### Nuclear localization of YAP leads to activation of the AT1 program in iAT2s

In order to examine the roles of YAP independent of upstream Hippo pathway inputs, we engineered a lentiviral vector encoding a constitutively active nuclear YAP cassette, YAP5SA, in which 5 serine residues have been mutated to alanines, preventing phosphorylation by LATS kinases^51^ (Fig. 2a). We differentiated human iPSCs into iAT2s using our previously established protocol^29, 52^ (Fig. 2b), and transduced the resulting iAT2s (>95% SFTPC^tdTomato^+) with lentiviral YAP5SA vs. parallel untransduced controls (mock). After 12 days of outgrowth in 3D cultures maintained in our published iAT2 media (CK+DCI),^29^ we observed significant upregulation of YAP target genes, *CTGF, ANKRD1*, and *CYR61*, in the YAP5SA transduced samples (Fig. 2c). YAP5SA overexpression resulted in a significant decrease in the AT2 markers *SFTPC* and *NAPSA*, and a significant increase in the 4-gene AT1 marker set, *AGER, CAV1, PDPN*, and *CLIC5* (Fig. 2c). While lung epithelial marker NKX2-1 was slightly decreased, we observed little to no expression of mesenchymal (*SNAIL1* and *TWIST*), airway (*P63, SCGB3A2*), or non-lung endoderm (*TFF1, AFP, CDX2*) markers after lentiviral transduction (Fig S2a). These results suggest that nuclear YAP activity promotes loss of the AT2 program and gain of AT1 markers in iAT2s.

**Figure 2:**
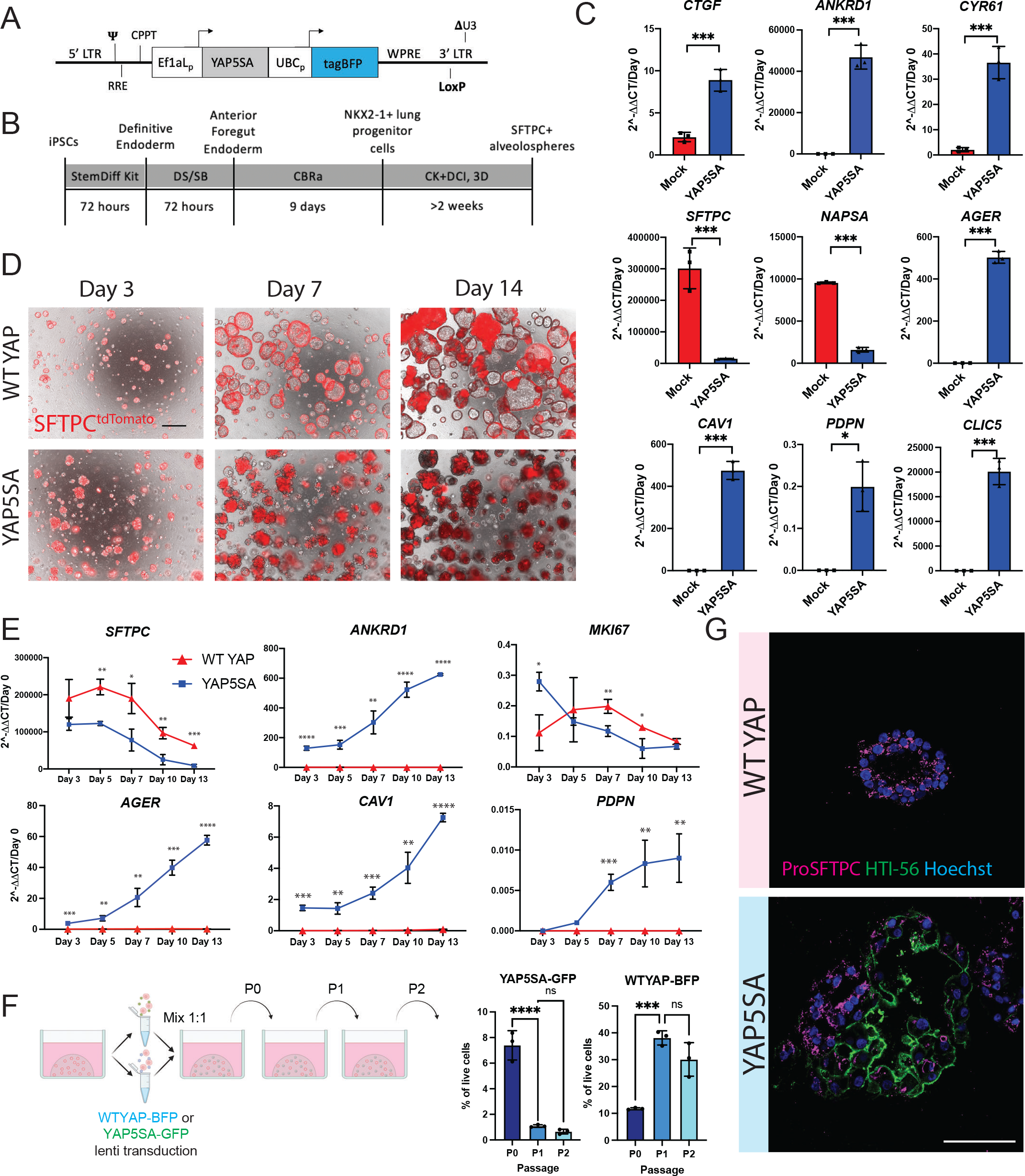
iAT2s transduced with nuclear YAP overexpression lentivirus upregulate AT1 marker genes. A) Diagram of lentiviral vector encoding dual promoters driving activated nuclear-specific mutated YAP (YAP5SA), the blue fluorophore tagBFP, and LoxP site. B) Directed differentiation protocol for producing iAT2s as previously published.^52^ (Stem Cell Technologies StemDiff endoderm kit, “DS/SB” = Dorsomorphin and SB43152, CBRa = Chir, BMP4, Retinoic Acid, CK + DCI = Chir, rhKGF, Dexamethasone, Cyclic AMP, IBMX, 3D plating in Growth Factor Reduced Matrigel.) C) Expression of indicated genes by RT-qPCR relative to Day 0 iPSCs in whole well RNA extracts taken 14 days post YAP5SA or mock lentiviral transduction of SPC2-ST-B2 iAT2s (>90% SFTPC^tdTomato^+ prior to lentiviral transduction, N=3 transductions. Student’s t test). D) Representative live cell imaging of SPC2-ST-B2 iAT2s following transduction with either WT YAP or YAP5SA lentivirus. (Brightfield/SFTPC^tdTomato^ overlay, scale bar = 500um). E) Whole-well gene expression by RT-qPCR over time following WT YAP vs. YAP5SA lentiviral transduction of SPC2- ST-B2 iAT2s, relative to Day 0 iPSCs. N=3 transductions. F) Competition assay – SPC2-ST-B2 iAT2s were transduced with either a WT YAP–tagBFP or YAP5SA-GFP lentivirus and mixed 1:1 before 3D plating. Cells were assessed by FACS and passaged normally every 14 days for 3 passages (1-way ANOVA). G) Immunofluorescence staining for ProSFTPC (magenta) and HTI- 56 (green) (Nuclei, blue; Scale bar = 100um). *p<0.05, **p<0.01, ***p<0.001, and ****p<0.001 for all panels.

To better control for lentiviral forced overexpression of the YAP transcript, we generated a control lentivirus with the same construct encoding constitutively over-expressed wild type (WT) YAP (Fig. S2b), which is predicted to be phosphorylated and degraded with minimal nuclear translocation.^51^ iAT2s transduced with WT YAP continued to grow normally as monolayered epithelial spheres with visible lumens and retained SFTPC^tdTomato^ reporter expression, whereas YAP5SA transduced cells rapidly lost SFTPC^tdTomato^, and by 3 days post-transduction exhibited an altered morphology with aggregated clumps of cells lacking visible lumens (Fig 2d). These morphological differences became more pronounced over time, as documented by microscopy at both 7- and 14-days post transduction (Fig 2d).

We hypothesized that activated nuclear YAP might be driving a transition in iAT2 transcriptomic programs from AT2-like to AT1-like. To further study the kinetics of this hypothesized transcriptomic shift following YAP5SA lentiviral transduction, we performed a time series gene expression analysis by taking whole well RNA extracts over a 2-week period post infection with lentiviral YAP5SA vs WT YAP. By day 3 after infection, *SFTPC* expression was already decreased in the YAP5SA infected well compared to WT YAP control and continued to decrease over time. There was some decline in *SFTPC* in WT YAP cells with time, which was consistent with our prior observations following iAT2s in culture as they approach confluence, prior to further passaging.^53^ AT1 markers *AGER, CAV1* and *PDPN* were significantly upregulated and continued to increase over time with only PDPN seeming to plateau by 2 weeks. The YAP downstream target *ANKRD1* followed a similarly increasing pattern (Fig 2e).

Consistent with the known non-proliferative, quiescent state that characterizes AT1s in vivo,^54^ decreased proliferation in outgrowth cells was suggested by: 1) decreasing expression of *MKI67* over time post YAP5SA transduction (Fig 2e), and 2) loss of YAP5SA infected cells after serial passaging using a competition assay between iAT2s transduced with WT YAP-BFP vs. YAP5SA- GFP lentiviruses (Fig 2f, S2b). Emergence of the AT1 program in YAP5SA transduced cells in 3D culture was also evident at the protein level by immunostaining for the AT1 marker, HTI-56 (Fig 2g). Whereas iAT2s exposed to the WT YAP lentivirus gave rise to epithelial spheres expressing only punctate proSFTPC cytoplasmic protein with no detectable HTI-56 staining, iAT2s exposed to YAP5SA lentivirus gave rise to subsets of organoids that exclusively contained either HTI-56 positive cells, or proSFTPC-positive cells, with other subsets of organoids containing a mixture of cells positive for either HTI-56 or proSFTPC (Fig. 2g).

### YAP5SA-induced acquisition of the iAT1 program profiled by single cell RNA sequencing

To further evaluate the effect of nuclear YAP on the global transcriptomic programs of iPSC-derived alveolar epithelial cells and to understand the heterogeneity of the lentiviral transduced cells at single cell resolution, we repeated YAP5SA vs WT YAP lentiviral transduction experiments and performed profiling by scRNA-seq (SPC2 iPSC line; SPC2-ST-B2 clone^55^; Fig 3a). We analyzed the resulting cells at 7 days post transduction, the time point at which there were significant differences in all genes measured by bulk RT-qPCR (Fig 2e). All live cells were captured for these profiles to allow examination of both transduced as well as non-transduced cells within the same well.

**Figure 3:**
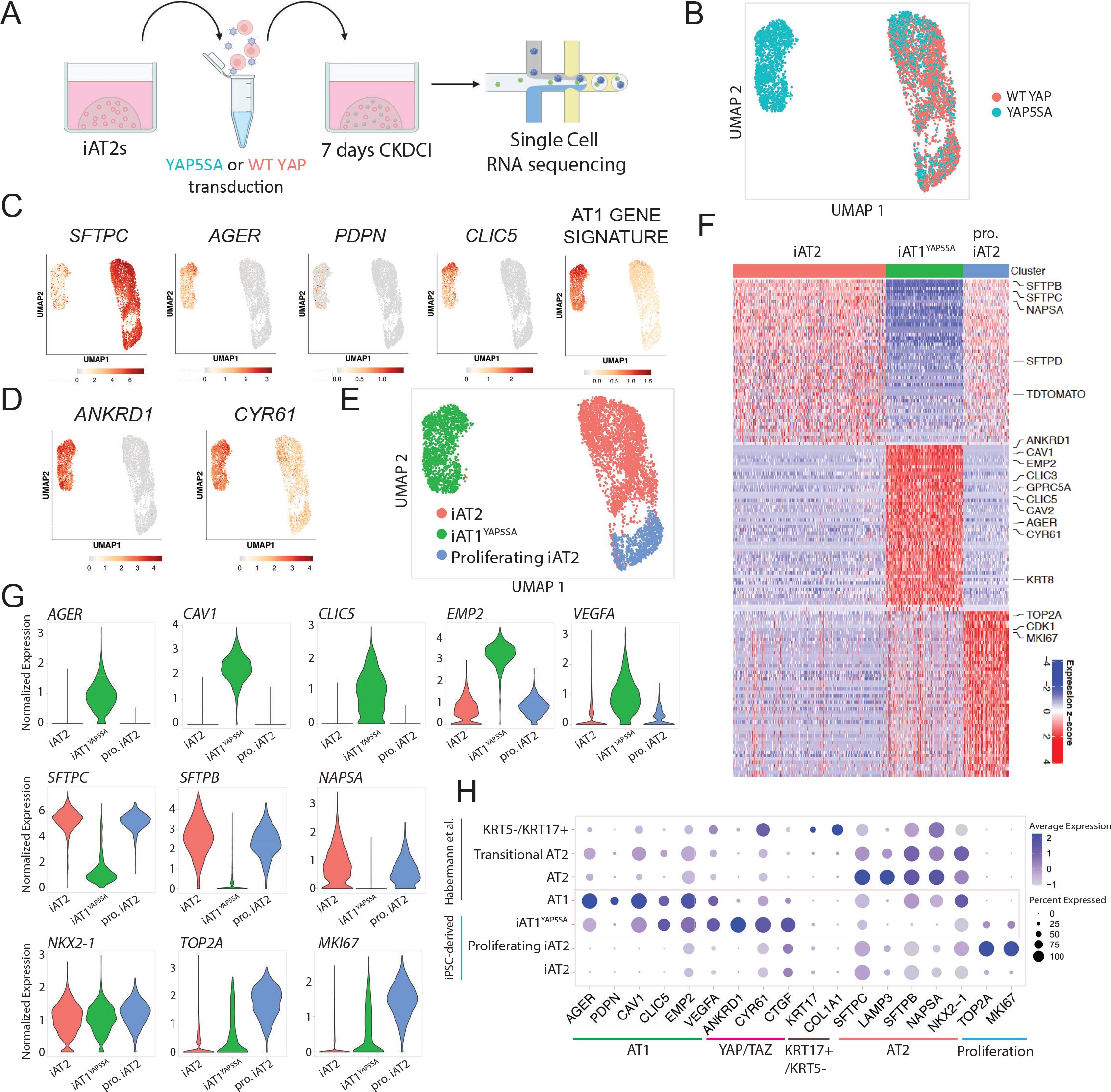
Nuclear YAP overexpression drives a shift from AT2 to AT1 programs in a cell autonomous manner. A) Experimental schematic for single cell RNA sequencing of WT YAP and YAP5SA lentiviral transduced iAT2s 7 days post transduction. B) UMAP of transcriptomes profiled from WT YAP and YAP5SA exposed samples. C) Gene expression overlays of: AT2 marker *SFTPC*, AT1 markers *AGER, PDPN, CLIC5*, a 50 gene AT1 signature figure 1 (Supplemental Table 1), and D) YAP downstream target genes *ANKRD1* and *CYR61*. E) Louvain clustering at resolution 0.05 identifies 3 indicated clusters. F) Top 50 differentially upregulated genes for the clusters shown in E. G) Gene expression of specific AT2, AT1, and proliferation markers across the clusters shown in E (“pro. iAT2”= proliferating iAT2s). H) Dot plot showing expression levels and frequencies of the indicated genes in iPSC-derived cells compared to scRNA-seq profiles of human adult primary AT1, AT2, Transitional AT2 and KRT5-/KRT17+ populations previously published by Habermann et al.^2^

By visualization of populations in UMAP space, the WT YAP-transduced control iAT2 population clustered together regardless of lentiviral transduction as shown by lentiviral BFP expression (Fig 3b, Fig S3a). This population showed high expression of the AT2 marker *SFTPC* (Fig 3c), as expected. The population of cells exposed to YAP5SA lentivirus contained some cells clustered with the WT YAP cells and some that clustered separately (Fig. 3b), presumably resulting from heterogeneity in transduction efficiency. Consistent with this speculation, expression of the BFP reporter present in our lentiviral construct localized mostly to the newly emergent cell cluster (Fig S3a). To screen for emergence of the AT1 program, we analyzed expression of AT1 markers *AGER, PDPN*, and *CLIC5* as well as our AT1 50 gene-set signature derived from the primary human lung dataset in figure 1 (Fig 3c-g, Table S1, Fig S3b). We observed high *AGER* and AT1 gene expression in the new cluster made only from the YAP5SA-transduced cells with little to no expression in the WT YAP transduced cluster. This upregulation of the AT1 program coincided with high expression of YAP downstream targets *ANKRD1* and *CYR61* (Fig 3d), and loss of expression of AT2 markers such as *SFTPC, SFTPB*, and *NAPSA* (Fig 3c-g).

We employed unbiased Louvain clustering to identify three distinct cell clusters (Fig. 3e), which we annotated based on the top 50 DEGs enriched in each cluster (Fig. 3f and S3c; Table S2). Both iAT2s and proliferating iAT2s expressed similarly high levels of AT2 marker genes such as *SFTPC, SFTPB*, and *NAPSA*, whereas proliferating iAT2s were uniquely enriched in expression of proliferation markers such as *MKI67* and *TOP2A* (Fig. 3f, g). The YAP5SA driven cluster’s top differentially expressed genes included YAP downstream targets *ANKRD1* and *CYR61* as well as AT1 marker genes such as *AGER, CAV1, CAV2, CLIC3, CLIC5, VEGFA*, and *EMP2* (Fig. 3f, g). Known markers for transitional or aberrant KRT17+/KRT5- AT2-derived cells^2^ were not enriched at significant levels in our YAP5SA transduced cells, though detectable expression of some of these markers was seen in both iAT2 and iAT1^YAP5SA^ populations (Fig S3d). Markers of airway epithelia (*SCGB3A2, FOXJ1, TP63*) were expressed very rarely, if at all (Fig S3d).

To provide an unbiased assessment of the lineage identity of our YAP5SA induced cell cluster, hereafter iAT1^YAP5SA^, based on the top differentially expressed genes enriched in this cluster (FDR<0.05, logFC>1; 92 genes), we employed the Tabula Sapiens gene set library,^56^ and found “Lung- type I pneumocyte” to be the top enriched term in our cluster (ranked by either p value or odds ratio), compared to all profiled tissue lineages (Fig S3e). Additional comparisons to primary human lung scRNA-seq datasets published by Habermann et al.^38^ indicated similar expression frequencies of individual AT1 markers in iAT1^YAP5SA^ cells compared to primary AT1s in vivo (Fig. 3h). However, iAT1s had higher frequency of expression of YAP downstream targets, *ANKRD1*, *CYR61,* and *CTGF*, consistent with forced over-expression of activated nuclear YAP (Fig. 3h). Taken together, our results suggest that overexpression of nuclear YAP drives the downregulation of the AT2 program and upregulation of the AT1 program in human iAT2s. Since cells grown in the same well but not transduced with the YAP5SA lentivirus maintained their iAT2 identity and clustered with the WT YAP control cells, this suggests nuclear YAP acts in a cell-autonomous manner to initiate this differentiation program without evidence of paracrine effects on other cells.

### Cre Excision of YAP5SA lentivirus results in reversion to the iAT2 state

We next sought to determine whether iAT1^YAP5SA^ cells stably maintain this cell program after removal of forced over-expression of YAP5SA. Recent publications indicate that YAP/TAZ knock out in murine AT1s leads to a loss of AT1 markers and expression of AT2 markers suggesting a reversion to AT2 cells.^57, 58^ Since we engineered the lentiviral YAP5SA vector to carry a loxP site in the lentiviral 3’LTR which is copied to the 5’LTR after infection, resulting in a floxed vector after genomic integration (Fig 2a, S2b), we employed adenoviral Cre-mediated vector excision (AdenoCre) and assayed the phenotype of the resulting cells after removal of YAP5SA (Fig S4a). By fluorescence microscopy and flow cytometry, AdenoCre infected cells augmented SFTPC^tdTomato^ expression compared to uninfected YAP5SA cells (Fig S4b,c). Additionally, YAP5SA-AdenoCre infected cells resumed proliferation at rates comparable to WT YAP spheres as quantified by EdU incorporation (Fig S4c). However, cell counts at this stage were not significantly different between conditions, possibly due to cell loss or toxicity from adenoviral infection as suggested by the WT YAP Adeno-Cre control sample (Fig S4d). YAP5SA-AdenoCre treated cells decreased expression of YAP downstream targets *CTGF* and *ANKRD1*, although not to levels of WT YAP suggesting only partial Cre excision. YAP5SA-AdenoCre cells also decreased expression of AT1 markers *AGER* and *CAV1* (although still present) and increased *SFTPC* expression, reverting to levels similar to WT YAP control iAT2s (Fig S4e). Together, these data suggest that AT1-like cells generated through forced activation of nuclear YAP signaling do not exhibit a stable AT1 phenotype, at least when maintained in CK+DCI medium, a condition that has been optimized for iAT2 maintenance.

### Development of AGER reporter iPSCs to track iAT1s

We next sought to engineer a fluorescent reporter to identify, monitor, and purify human iAT1s. Due to its high expression in and specificity to AT1 cells in our human primary single cell dataset (Fig 1f), *AGER* was selected as a candidate human AT1 marker locus for reporter targeting. Our lab has previously published the use of gene editing to create an NKX2-1^GFP^ reporter iPSC line (BU3 NG; Hawkins et al. 2017)^59^ for the visualization and purification of NKX2-1+ lung epithelial cells. Using CRISPR/Cas9 targeting of iPSCs, we inserted a second fluorophore-encoding cassette, tdTomato, into the start codon of the endogenous human *AGER* locus of this same line (Fig. 4a, Fig S5a, b), generating a karyotypically normal bifluorescent reporter line carrying NKX2- 1^GFP^ and AGER^tdTomato^, hereafter BU3 “NGAT”.

**Figure 4:**
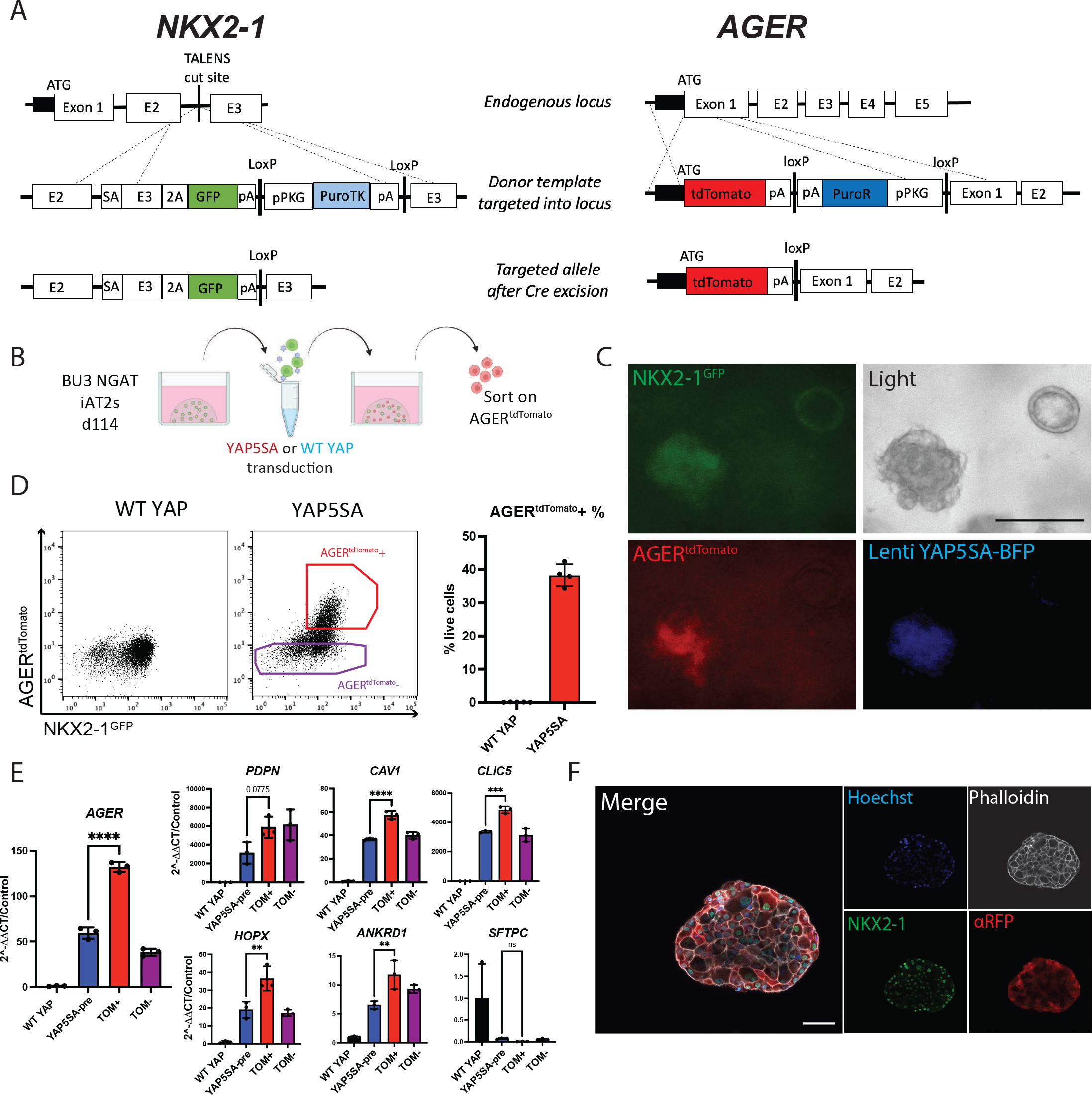
A bifluorescent NKX2-1^GFP^; AGER^tdTomato^ reporter iPSC line enables tracking and purification of iAT1^YAP5SA^ cells. A) Gene editing strategy to generate the BU3 NKX2-1^GFP^/AGER^tdTomato^ (NGAT) dual reporter iPSC line for tracking lung epithelial lineages and AT1-like cells. (BU3 NKX2-1^GFP^ reporter previously published by Hawkins et al.)^59^ B) BU3 NGAT iPSCs were differentiated into iAT2s and transduced with WT YAP or YAP5SA lentivirus. AGER^tdTomato^+ cells, appearing only in the YAP5SA well, were then sorted and analyzed after 14 days of outgrowth in CK+DCI. C) Live cell fluorescence microscopy of YAP5SA-transduced cells growing next to an un-transduced epithelial sphere showing GFP, tdTomato, and TagBFP fluorescence; Scale bar = 200um). D) Flow cytometry analysis of iAT2s (BU3 NGAT line) 14 days post WT YAP or YAP5SA lentiviral transduction showing sorting gate for AGER^tdTomato^ + and – cells with AGER^tdTomato^+ percentage quantified. E) Gene expression analysis of sorted cells from D, compared to WT YAP and YAP5SA unsorted cells by RT-qPCR. WT YAP = Unsorted WT YAP transduced cells, YAP5SA-pre = Unsorted YAP5SA transduced cells, TOM+ = AGER^tdTomato^+ sorted cells, TOM- = AGER^tdTomato^- sorted cells. (1-way ANOVA). F) Immunofluorescence staining of NKX2-1 protein and tdTomato (*α*RFP) (NKX2-1: green, F-actin: Phalloidin white, tdTomato: red, Nuclei: blue. Scale bar = 50um). *p<0.05, **p<0.01, ***p<0.001, and ****p<0.001 for all panels.

We characterized BU3 NGAT iPSCs after directed differentiation into distal lung epithelium followed by YAP5SA lentiviral transduction (Fig. 4b). Cells transduced with the WT YAP lentivirus did not express tdTomato, while YAP5SA transduced cells expressed AGER^tdTomato^ specifically in the NKX2-1^GFP^+ lung epithelial population (Fig 4c, d). Additionally, the parental BU3 NG line, when transduced with YAP5SA, showed no tdTomato expression (Fig S5c). BU3 NGAT-derived AGER^tdTomato^+ cells were sorted and their expression levels of YAP downstream targets, AT2 markers, and AT1 marker genes were compared to that of YAP5SA-transduced unsorted (“presort”) cells, tdTomato- sorted cells, or control unsorted cells from WT YAP transduced samples (Fig 4e). AGER^tdTomato^+ cells were enriched in expression of *AGER* as well as AT1 markers *CAV1, PDPN*, and *CLIC5* (Fig 4e), suggesting the utility of the reporter to track and purify iAT1s. Immunofluorescence microscopy further confirmed tdTomato protein expression was specific to cells co-expressing nuclear NKX2-1 protein, and the altered clumped organoid morphology observed in YAP5SA transduced SPC2-ST-B2 iAT2s was recapitulated in transduced cells derived from the BU3 NGAT line (Figs 2, 4f).

### Serum free medium-based induction of iAT1s

We next sought to identify factors that could be added to a defined medium to recapitulate iAT2 to iAT1 differentiation without requiring lentiviral forced YAP over-expression. Genetic mouse models of transgenic over-expression of activated YAP or conditional deletion of the Hippo kinases LATS1 and LATS2 in the developing lung epithelium have been shown to increase expression of mouse AT1 markers, suggesting a role for Hippo-LATS-YAP signaling in AT1 differentiation.^48^ More recently, the small molecule LATS inhibitor LATS-IN-1 (hereafter “L”) has been shown to drive nuclear YAP *in vitro*.^60^ Thus, we tested the effect of L supplementation on iAT2s cultured with and without the iAT2 supportive growth factors CHIR99021 (Chir, or “C”) and KGF/FGF7 (“K”) in our base media (“DCI”) (Fig 5a-e). We found L induced rapid and robust changes in iAT2s including: 1) morphologic changes within 48 hours, resembling those of the YAP5SA-transduced cells, and 2) AGER^tdTomato^ induction, detectable within 72 hours in the NKX2-1+ population and increasing over the following 2 weeks to 84.97±1.23% of the NKX2-1^GFP+^ cells in L+DCI (Fig. 5b-d). Comparing L supplemented conditions with/without Chir and KGF, the AGER^tdTomato^ expression was highest and most frequent in wells without Chir or KGF, suggesting inhibitory effects of these growth factors (Fig 5b, d, e). There was little to no AGER^tdTomato^ expression observed in cells exposed to DCI without L (Fig 5b, d, e).

**Figure 5:**
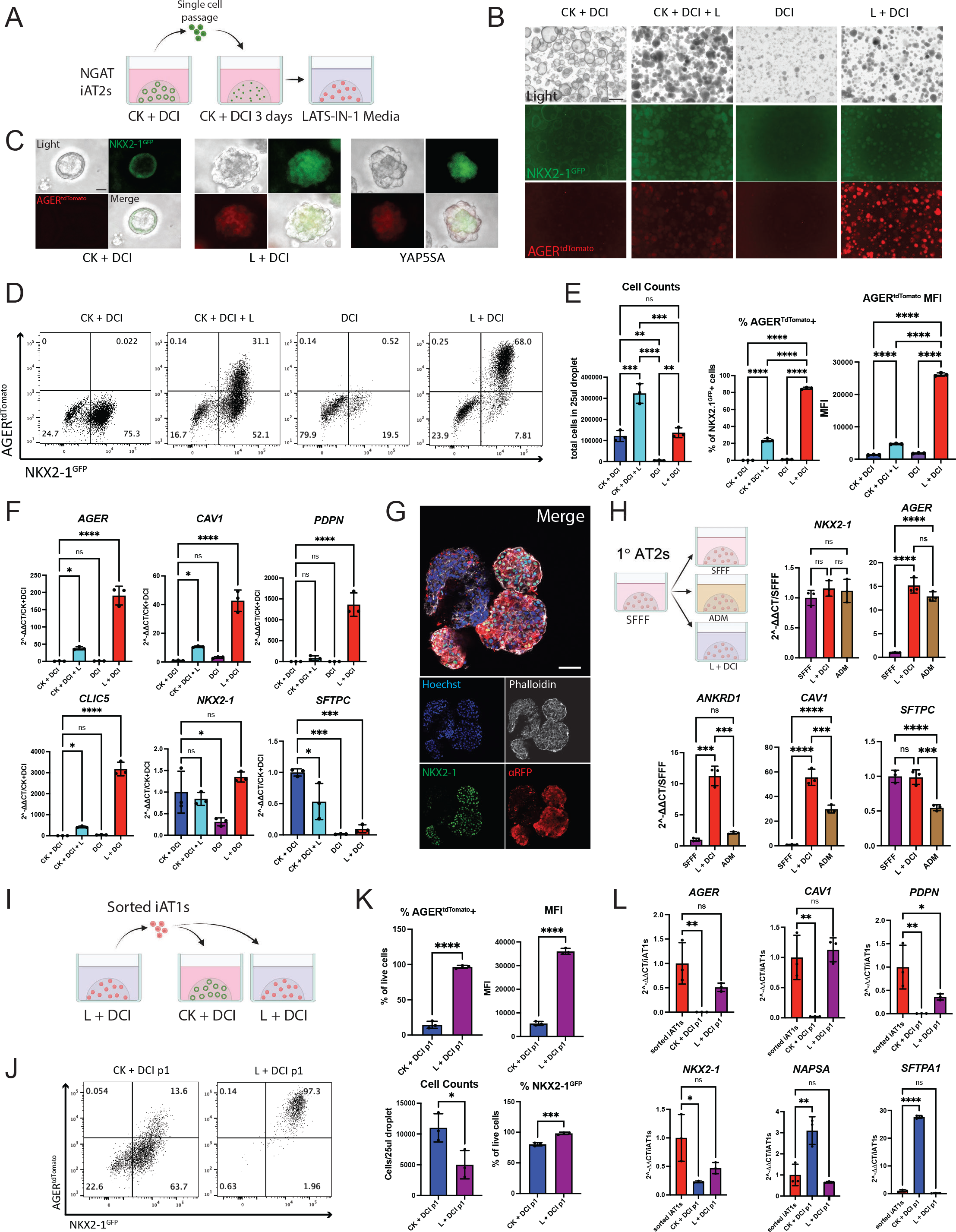
Serum-free medium-based induction of AGER^tdTomato^ A) BU3 NGAT iAT2s were passaged as usual into 3D Matrigel with CK+DCI + RI medium for 3 days. Medium was then kept the same or switched to LATS inhibitor-based media, CK DCI + L, DCI, or L DCI. (C = Chir, K = rhKGF, L = LATS-IN-1; “DCI” as defined in figure 2). B) Representative live fluorescence microscopy of iAT2s (BU3 NGAT clone) at 9 days after changing to the indicated medium, either without (B) or with (C) YAP5SA transduction (Scale bars = 500 um in B; 50um in C). D) Representative flow cytometry plots of NKX2-1^GFP^ and AGER^tdTomato^ 9 days post after changing to each indicate medium. E) Cell counts and flow cytometry quantification of AGER ^tdTomato^+ percentage and Mean Fluorescence Intensity (MFI) 9 days post medium change. One-way ANOVA, N=3 per condition. F) Gene expression of AT1 and AT2 markers by whole well RT-qPCR. (1-way ANOVA, N=3 per condition). G) Whole mount immunofluorescence microscopy of BU3 NGAT organoids in L+DCI. (NKX2-1: green, F-actin: Phalloidin white, tdTomato: red, Nuclei: blue. Scale bar = 50um.) H) Primary adult human AT2 cells (1°AT2s) were cultured as described^32^ and 7 days post plating, medium was changed to published, human-serum containing Alveolar Differentiation Media (ADM) or L+DCI. Whole well RT-qPCR of selected genes 7 days post medium change. 1-way ANOVA, N=3 per condition. I)Experimental schematic: BU3 NGAT iAT1s were grown in L+DCI for 10 days, sorted on AGER^tdTomato^, and then plated in 3D Matrigel in either L+DCI or CK+DCI. Outgrowths of sorted cells were analyzed after an additional 9 days. J) Representative flow cytometry plots of AGER^tdTomato^+ outgrowth from (I). K) Quantification of indicated fluorescence (FACS), cell counts, or transcript expression (L; whole well RT-qPCR) of cells from (I). Control sorted iAT1s in L are freshly sorted AGER^tdTomato^+ cells after 9 days in L+DCI. (N=3 per condition; K=Student’s t test; L=1-way ANOVA). *p<0.05, **p<0.01, ***p<0.001, and ****p<0.001 for all panels.

We then examined the individual effects of Chir and KGF in inhibiting AGER^tdTomato^ in media containing L and saw that both were significantly inhibitory to both AGER^tdTomato^ percentage and mean fluorescence intensity (MFI) (Fig S6a, b). To test whether this was specific to the LATS inhibitor media-based differentiation, we also tested withdrawal of each of these factors in our YAP5SA differentiation model. We saw inhibitory effects of both growth factors on AGER^tdTomato^ as well as AT1 markers by bulk RT-qPCR, suggesting both Chir and KGF need to be removed for the efficient differentiation of iAT1s (Fig S6c-g). This is consistent with prior publications suggesting canonical Wnt or KGF signaling are inhibitory to expression of the AT1 program.^20, 26, 34^

After dose response testing to identify an optimal dose of L, replacing the CK+DCI medium on day 3 after passaging iAT2s with a medium consisting of L+DCI using a 10uM dose of LATS-IN- 1, yielded the greatest frequency and brightness (MFI) of NKX2-1^GFP^/AGER^tdTomato^+ cells 2 weeks post passage (Fig S7a). When NKX2-1^GFP+^ cells were re-sorted to purify the lung epithelial population and exposed to L+DCI beginning 3 days post sort, up to 97% of all cells were AGER^tdTomato^+ by day 14 (Fig S7b). By whole well RT-qPCR, *AGER, CAV1, PDPN*, and *CLIC5* were significantly upregulated in wells containing the LATS inhibitor, consistent with induction of a broad AT1 program, with higher levels expressed in conditions without KGF and Chir (Fig 5f and S7c). NKX2-1 remained unchanged between iAT2 media and media containing LATS-IN-1, suggesting maintenance of the lung epithelial program, whereas SFTPC was decreased (though not entirely absent) indicating downregulation of the AT2 program (Fig 5f and S7c). Similar trends were seen in SPC2-ST-B2 iAT2s grown in L+DCI (Fig S7d-f). Immunostaining showed similar organoid shape to that of YAP5SA-transduced organoids, with AGER^tdTomato^ only being expressed in NKX2-1+ cells (Fig 5g). Taken together these results indicate that treatment with the LATS inhibitor, L, in the absence of Chir and KGF efficiently differentiates iAT2s into iAT1s.

To determine whether L+DCI medium similarly induces the AT1 program in primary adult human AT2s, we first culture-expanded primary AT2s in serum-free, feeder-free conditions (SFFF) as reported.^32^ After seven days in SFFF, medium was either continued as SFFF or switched to published human serum-containing Alveolar Differentiation Medium (ADM) or L+DCI. Seven days later, *NKX2-1* expression was similar for all conditions, while *AGER* was significantly upregulated in both L+DCI and ADM conditions and was not significantly different between the two media. *CAV1* and *ANKRD1* expression was higher in L+DCI than ADM, suggesting this differentiation medium is effective in the differentiation of primary human AT2s into AT1s (Fig 5h). In tests using iAT2s instead of primary AT2s, only L+DCI and not ADM promoted iAT1 differentiation based on low induction of *CAV1* in response to ADM and no induction of *AGER* or *ANKRD1* (Fig S7g).

To determine whether iAT1s maintain their phenotype when transitioned back into iAT2- maintenance medium (CK+DCI), AGER^tdTomato^+ cells were sorted after 11 days in L+DCI and replated into 3D Matrigel in either L+DCI or CK+DCI (Fig 5i). After 9 further days in culture, cells in L+DCI maintained >95% AGER^tdTomato^ expression, while those in CK+DCI showed 10-20% AGER^tdTomato^+ cells (Fig 5i-l) and re-expressed AT2 markers consistent with residual plasticity in iAT1s. Those iAT1s maintained in L+DCI, compared to CK+DCI, better maintained the AT1 program as evidenced by retained expression of *AGER* and *CAV1* and suppression of AT2 markers (Fig 5j-l).

#### Transcriptomic profiling by scRNA-seq of iAT1s

To comprehensively compare the transcriptomic programs of iAT1s generated through our defined iAT1 medium (L+DCI) vs our method of lentiviral forced over-expression of YAP5SA in CK+DCI, we performed head-to-head profiling of iAT1s prepared by either method vs parallel controls maintained as iAT2s, hereafter named iAT1, iAT1^YAP5SA^, and iAT2 respectively (SPC2- ST-B2 iPSCs; Fig 6a). Clustering analysis revealed 3 predominant clusters (Louvain resolution =0.05; Fig 6a, b) visualized by UMAP which segregated based on the method used to generate the cells. As in figure 3, cells from the YAP5SA sample segregated into either the iAT1^YAP5SA^ cluster or the iAT2 cluster, depending on whether they were successfully transduced (iAT1^YAP5SA^) or not transduced (iAT2s) with the lentiviral vector (see inset figure 6a).

**Figure 6:**
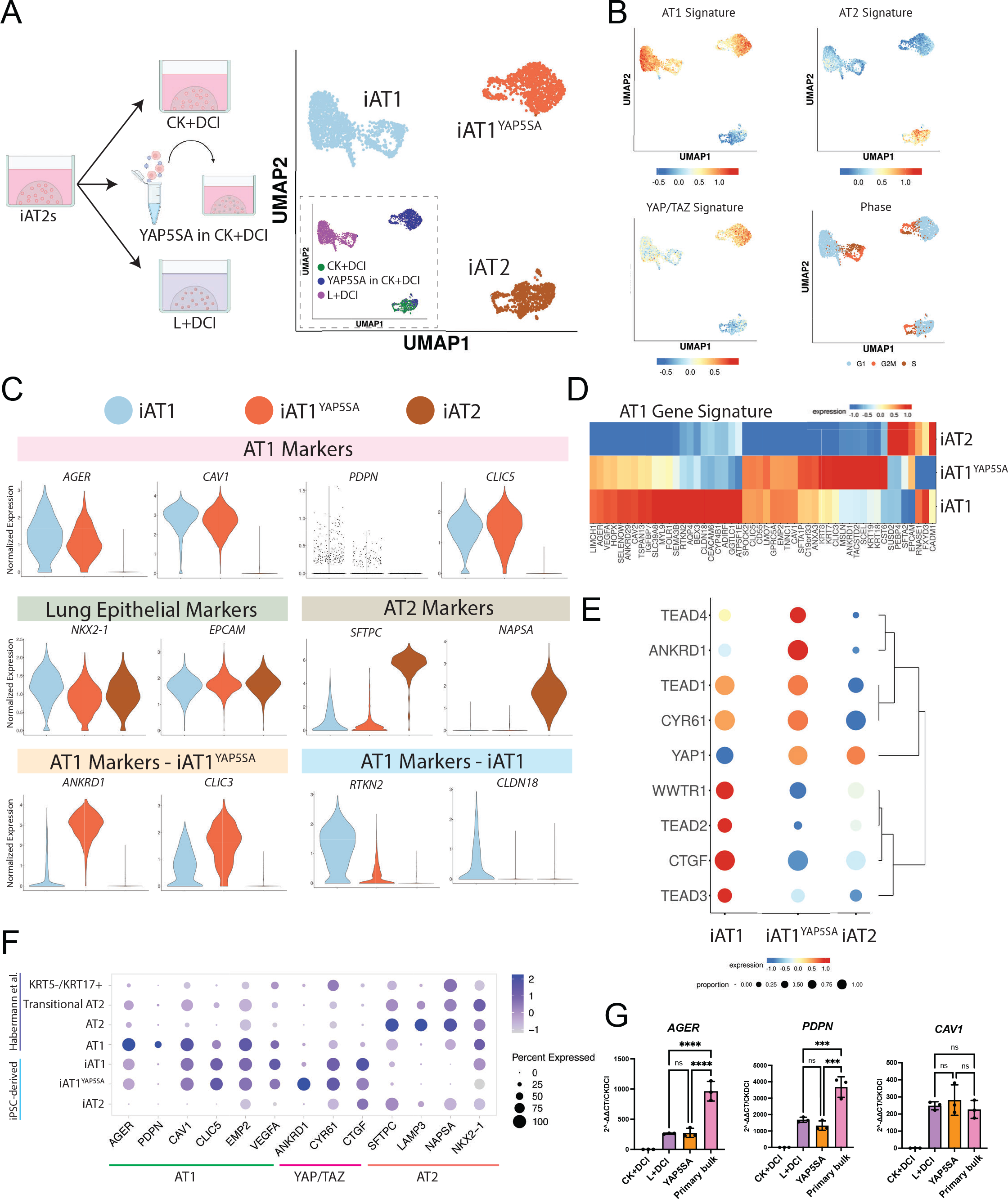
iAT1s generated by either a defined medium or by lentiviral activated nuclear YAP express a broad AT1 transcriptomic program. A) Experimental schematic (left panel) showing SPC2-ST-B2 iAT2s were either grown in CK+DCI, transduced with YAP5SA lentivirus and grown in CK+DCI, or grown in CK+DCI for 3 days before switching to L+DCI. Nine days post passage live cells were profiled by scRNA-seq. UMAP visualization (right panel) of single cell transcriptomes by either original sample name (inset) or after Louvain clustering at resolution 0.05 into 3 clusters named iAT2, iAT1^YAP5SA^, and iAT1. B) Gene expression overlays of a human AT1 50-gene signature, a 22 gene YAP/TAZ signature,^61^ human AT2 50-gene signature, or cell cycle phase. C) Violin plots quantifying expression of indicated markers across the clusters from (A). D)Heatmap showing average expression (normalized by column) of genes in the AT1 50-gene set across the 3 Louvain clusters from (A). E) Dot plot of transcript expression levels and frequencies of YAP/TAZ downstream targets and TEADs across the 3 clusters from (A). F) Dot plot showing expression levels and frequencies of AT1, AT2, and YAP/TAZ targets in this dataset compared to human adult primary AT1, AT2, Transitional AT2, and KRT5-/KRT17+ scRNA-seq profiles previously published by Habermann et al.^2^ G) Expression of AT1 marker genes (RT-qPCR) in whole well RNA extracts of L+DCI-induced and YAP5SA-transduced iAT1s compared to bulk primary human distal lung tissue; fold change normalized to 18S (2^-DDCT) is calculated relative to iAT2s in CK+DCI. (1-way ANOVA) *p<0.05, **p<0.01, ***p<0.001, and ****p<0.001 for all panels.

We observed that both iAT1 and iAT1^YAP5SA^ populations exhibited downregulation of the AT2 gene signature with frequent and robust upregulation of the primary AT1 50-gene set signature (Fig 6b, Table S1) as well as individual AT1 marker genes (*AGER, CAV1, PDPN*, and *CLIC5*; Fig 6c-g, Fig S81, b), consistent with differentiation of iAT2 into AT1-like cells using either method. Upregulation of AT1 transcripts, *AGER, PDPN*, and *CAV1* was validated by RT-qPCR (Fig 6g), and the frequency of expression of most AT1 marker transcripts was similar to expression profiles of published primary AT1s (Fig. 6f), although absolute expression levels for *AGER* and *PDPN* were lower than in primary control distal lung tissue by RT-qPCR (Fig 6g, S8c).

Despite these similarities, iAT1 and iAT1^YAP5SA^ cells clustered separately with 298 transcripts differentially expressed (FDR<0.05, logFC>1, Table S2). These differences include: 1) differential upregulation of a YAP/TAZ 22-gene target signature^61^ (Fig 6b), likely reflecting the higher levels of YAP signaling induced by forced over-expression of the YAP5SA driver compared to milder upregulation resulting from the LATS inhibitor, and 2) differential expression of other AT1 marker genes in our AT1 50 marker set, such as *ANKRD1* and *CLIC3* which were more highly expressed in the iAT1^YAP5SA^ cluster, or *RTKN2* and *CLDN18*, which were more highly expressed in the iAT1s (Fig 6c,d Fig S8a). Consistent with the above differences in the YAP/TAZ signature set were notable differences in specific Hippo-LATS-YAP signaling targets. For example, *YAP* and *TEAD4* were upregulated in iAT1^YAP5SA^ cells, whereas *TAZ (WWTR1), TEAD2, TEAD3*, and *CTGF* were upregulated in iAT1s (Fig 6e). While it is possible that some of these differences are due to the LATS inhibitor affecting both YAP and TAZ whereas the lentivirus is YAP-specific, they could also be due to the effect of Chir and KGF in the medium of the lentiviral-transduced iAT1^YAP5SA^ cells.

We found that both iAT1^YAP5SA^ and iAT1 populations included a subset of cells in active cycle (Fig 6b) and both contained minor subsets expressing detectable levels of some, but not all, transitional state markers, such as *KRT17* (Fig S8b). Importantly, neither population showed significant expression of nonlung endoderm, Epithelial-Mesenchymal Transition (EMT), or airway (*FOXJ1, TP63*, or *SCGB1A1*) markers, except for *SCGB3A2* which was differentially upregulated in iAT1s (Fig S8b). A small subpopulation appeared in the iAT1 population that is high in expression of Notch signaling targets such as *HES1* but is lower in AT1 gene signature. Taken together, these results indicate efficient induction of the AT1 program without the need for lentiviral transduction via a serum-free differentiation medium, L+DCI.

#### iAT1s display functional capacities to form a flattened epithelial barrier, secrete signaling ligands, and express extracellular matrix-encoding transcripts

We next evaluated the functional capacity of iAT1s generated in defined L+DCI medium. An emerging literature has established several functions that characterize primary AT1s in vivo, including their potential to: form flattened cells that contribute to an epithelial barrier;^1^ produce a characteristic alveolar extracellular matrix;^62^ and serve as signaling hubs in the lung through secretion of ligands that bind receptors on adjacent lung mesenchymal lineages.^63^ Since iAT1s in 3D Matrigel cultures are still proliferative and do not discernably form flattened epithelial barriers, we transitioned BU3 NGAT iAT1s into transwell cultures in L+DCI, and 3 days later aspirated apical medium to establish air-liquid interface (ALI) cultures in L+DCI, hereafter “iAT1 ALI P1”. We observed maintenance of AGER^tdTomato^ reporter expression in the outgrowth cells at ALI culture, suggesting maintenance of the AT1 program (Fig 7a-c), comparable to 3D L+DCI culture conditions. We also noted formation of an epithelial barrier after ALI culture, based on prevention of apical medium leakage and increasing transepithelial resistance (TEER) over time to 1479±166Ω•cm^2^ by day 10 (Fig 7d).

**Figure 7:**
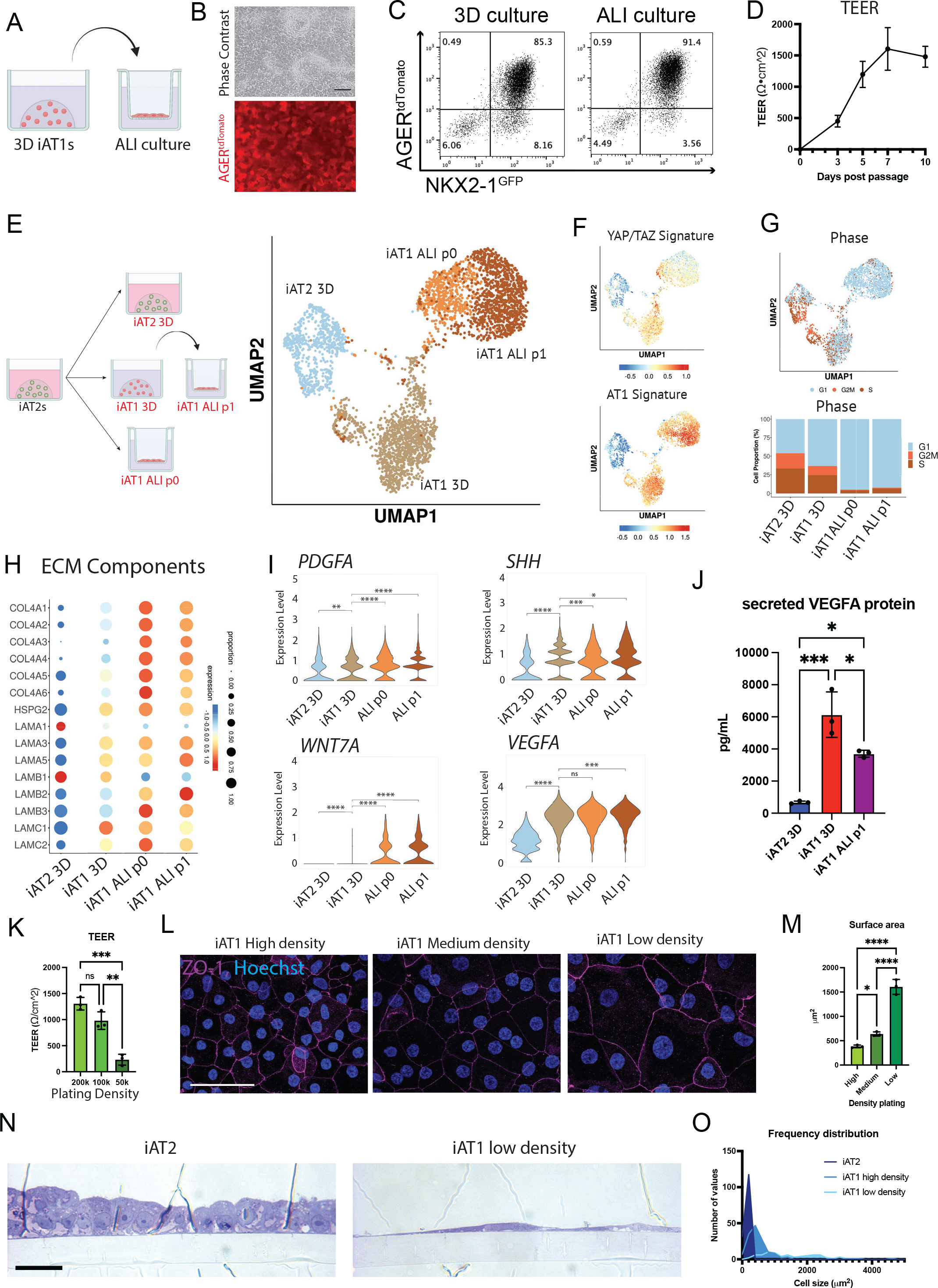
iAT1s cultured at air-liquid interface (ALI) express AT1-like molecular and functional phenotypes A) Experimental schematic indicating BU3 NGAT iAT1s were cultured in 3D in L+DCI medium for 8-11 days before single cell passaging and replating onto transwell inserts in L+DCI medium. Upper chamber media was aspirated after 3 days (airlift) to form an air-liquid interface (ALI). B) Live cell imaging showing retention of AGER^tdTomato^ in iAT1s after ALI culture. scale bar = 100um. C) Flow cytometric analysis of NKX2-1^GFP^ and AGER^tdTomato^ reporter expression in 3D or ALI cultures of iAT1s (6 days). D) Transepithelial electrical resistance (TEER) measurements of BU3 NGAT iAT1s over 10 days of ALI cultures (Air lifted at day 3; N=3). E) Profiling by scRNA-seq of BU3 NGAT iAT2s in 3D CK+DCI, iAT1s in 3D L+DCI, or iAT1s in ALI cultures. iAT1s in ALI cultures were plated either from iAT2s into L+DCI (iAT1 ALI P0) or were plated from 3D iAT1s after 9 days of pre-culturing in 3D L+DCI prior to transfer to ALI culture (iAT1 ALI P1). UMAP projection shows the 3D cultures cluster separately while the two iAT1 ALI cultures cluster closer together. F) UMAP overlays of YAP/TAZ 22 gene signature^61^ and primary human AT1 50-gene signature (Table S1). G) Cell cycle phase distribution across all samples. H) Expression of transcripts encoding extracellular matrix (ECM) components. I) Expression of transcripts encoding secreted ligands, comparing the samples from (E). J) Analysis of secreted VEGFA protein in conditioned media at day 10 of culture of each indicated sample. K) TEER of iAT1s after plating at High, Medium, and Low densities and outgrowth in ALI cultures in L+DCI. One-way ANOVA, N=3 per condition. L) Tight junction protein ZO-1 staining (magenta) at high, medium, and low plating density outgrowths at day 10 Scale bar = 50um. M) Average surface area of cells calculated using ZO-1 cell outlines at three different iAT1 ALI plating densities. N=3 per condition, averaged from ∼150 cells per sample. N) Cross sectional imaging of SPC2-ST-B2 iAT2s at ALI as previously published,^68^ and SPC2-ST-B2 iAT1s plated at low densities in L+DCI. (Toluidine Blue stain, scale bar = 10um). O) Frequency distribution of cell surface areas of iAT2, iAT1 high density, and iAT1 low density. N=149, 139, and 79 respectively). *p<0.05, **p<0.01, ***p<0.001, and ****p<0.001 for all panels.

We profiled the iAT1s produced in 3D L+DCI (iAT1 3D) vs iAT1 ALI P1 conditions as well as iAT1s produced in a second ALI condition where iAT2s are only exposed to L+DCI coincident with transwell plating (hereafter “iAT1 ALI P0”; Fig 7e-i, Fig S9a, b). Compared to parallel controls maintained as iAT2s, we found downregulation of the AT2 program and upregulation of the YAP/TAZ and AT1 programs in iAT1s grown in both ALI cultures, similar to those iAT1s produced in 3D-L+DCI, but with significantly reduced active cell cycling (reduced G2/M phases; Fig 7f-g, Fig S9a) and significantly less *TOP2A* (data not shown). These results indicate that iAT1s transition to a more quiescent state after either ALI culture. As we have previously reported for iAT2s,^64^ a rare subset of cells plated in these conditions upregulated markers of non-lung endoderm (*ALB, AFP*) suggesting some degree of residual plasticity in rare cells plated in all 4 conditions (Fig S9b).

Despite the similarities in expression levels of AT1 marker genes in cells cultured with vs without ALI conditions, iAT1s after ALI culture clustered separately from iAT1 3D, and genes encoding extracellular matrix components, such as *COL4A4*, featured prominently in those transcripts upregulated after ALI culture (Fig 7h; Table S2). Given the recent reported role of extracellular matrix generation by mouse AT1s during development,^62^ we looked at the other components of collagen IV, as well as laminins, including the components of laminin-332 (*LAMA3, LAMB3*, *LAMC2*). As reported for mouse AT1s in vivo,^62^ human iAT1s after ALI culture expressed all 6 components of collagen IV (with more frequent expression of *COL4A6* than the mouse), as well as patterns of laminin expression similar to published mouse single cell data (Fig 7h). Notably, in this mouse dataset, expression of the components of laminin-332 was associated with more mature AT1 cells, only turning on around E18 and continuing postnatally. However, the expression pattern of these laminins in human lung development is unknown.

One recently recognized function of AT1s in vivo is their role as signaling hubs in the distal lung, based on mouse and human scRNA-seq profiles and genetic mouse models that suggest AT1s stimulate local alveolar Shh, Wnt, PDGF, and VEGF signaling in adjacent alveolar mesenchymal or vascular endothelial lineages during alveologenesis or tissue maintenance.^62, 63^ Consistent with these reports, we found iAT1s after ALI culture upregulated transcripts encoding the signaling ligands, *PDGFA, SHH, WNT7A*, and *VEGFA* (Fig 7i) that were also enriched in primary AT1s from 2 month old human infants as reported by Zepp et al.^63^ At the protein level, we verified secreted VEGFA was present in conditioned medium from both 3D and ALI iAT1 culture conditions. (Fig 7j).

Finally, we evaluated whether iAT1s were capable of forming the characteristic flattened epithelial barrier that defines their unique morphology. We tested whether BU3 NGAT iAT1s would form a flattened monolayer at lower density plating, potentially stretching out to cover the available surface area, while still forming tight junctions necessary to form a functional epithelial barrier. Cells were plated at 3 separate densities in ALI cultures for 10 days. The ALI barrier remained intact for all 3 conditions with measurable TEER and formation of tight junctions, evident based on ZO-1 staining (Fig 7k,l). Average cell surface area was calculated, and cells plated at lower densities had significantly larger cell surface areas on average (Fig 7m, Fig S9c). Similar results were observed for SPC2-ST-B2 iAT1s plated at high- and low-density ALI cultures, and iAT1s occupied larger cell surface areas than iAT2s plated at an identical initial cell density (Fig 7n,o; S9d-f). Reported average surface areas of adult human AT1s range from 3960μm^2^ to 8290μm^2^, averaging around 5100μm^2^.^1, 5, 65^ At the above plating density, iAT1s reached the size of cells on the smaller end of this range (Fig 7o, S9c). Additionally, cross sectional images of SPC2-ST-B2 iAT1s show thin cells with elongated, flattened nuclei compared to the cuboidal iAT2s at ALI. These results suggest iAT1s readily flatten and stretch to cover available surface area in order to form and maintain a thin epithelial barrier.

## Discussion

In this study we employed directed differentiation of iPSCs in vitro to efficiently generate cells expressing several key features reminiscent of human AT1 cells. Using multiple human iPSC lines, we first derived iAT2s and found activated nuclear YAP signaling was sufficient to drive a global transcriptomic shift from the AT2 to the AT1 program. This transition could be recapitulated with a defined serum-free medium containing a LATS inhibitor, producing cells that broadly upregulated a YAP/TAZ signaling gene set, expressed the molecular phenotype of human AT1 cells, and exhibited AT1 functional features, including the potential to form a flattened epithelial barrier that expresses AT1-associated ligands and matrix components.

Our finding that Hippo-LATS-YAP signaling activates the human AT1 program in lung epithelia is consistent with several recent reports that employed mouse genetic models to activate this pathway in vivo, similarly finding resultant emergence of mouse AT1 markers.^23, 47–49, 58^ While nuclear YAP activation was sufficient to drive the transcriptomic shift from iAT2 to iAT1, withdrawal of canonical Wnt or FGF signaling activators (CHIR and KGF) alone was not sufficient to robustly induce this differentiation, as measured by AGER^tdTomato^ expression. We found these AT2- sustaining factors were inhibitory to the AT1 program, thus their withdrawal helped to promote the Hippo-LATS-YAP induced AT1 transition. In the 2 activating conditions we developed, we found that iAT1s grown in our defined medium (L+DCI) resembled YAP5SA-transduced iAT1s in terms of their structure, growth, and expression of most but not all canonical AT1 marker transcripts. While neither population was detectably more mature than the other, they clustered separately and differed in several ways with higher expression of the YAP target, *ANKRD1*, as well as expression of several keratins, *KRT18, KRT19*, and *KRT7*, all higher in the YAP5SA-transduced iAT1s, findings of unclear significance that will require further study. Importantly, our L+DCI medium, without requiring any lentivirus, also induced upregulation of several AT1 markers in cultured primary adult human AT2 cells, providing an identical serum-free defined medium-based approach that can now be studied using cells prepared from multiple sources, either primary or engineered.

Key to benchmarking our in vitro cells was the identification of gene set markers and pathways that define the in vivo transcriptomic program of human AT1 cells. Single cell RNA sequencing of adult primary human AT1 cells provided a comprehensive gene signature as well as a quartet of canonical markers (*AGER, PDPN, CAV1*, and *CLIC5*) that can be used for future profiling of these cells. Additionally, these datasets help to distinguish differences between human and mouse marker genes. While mice are the most common *in vivo* lung model, several mouse AT1 markers are not identically expressed in humans, including marker genes *HOPX* and *AQP5*, which are commonly used mouse AT1-specific genetic drivers that do not share this specificity in the human lung.^6, 40^ In prior mouse studies, AT1-selective fluorescent reporters or tracing approaches have revealed much about AT1 cell biology and lineage relationships.^6, 39, 40^ Similarly, we engineered a human AGER-targeted tdTomato fluorescent reporter iPSC line, providing a key reagent for the tracking, quantification, and purification of candidate human AT1-like cells. In combination with an NKX2-1^GFP^ reporter, we employed this bifluorescent system to understand the differentiation kinetics and characteristics of iAT1s as they emerge in various culture conditions from parental NKX2-1+ distal lung epithelial cells.

Additional mouse genetic models have shown that deletion of YAP/TAZ in mature AT1s in vivo leads to reversion to an AT2-like state.^22, 23^ Upon Cre-mediated excision of the YAP5SA lentivirus in our human iAT1^YAP5SA^ cells, we saw an upregulation of AT2 markers and an increase in iAT2 sphere morphology consistent with these mouse studies. Our medium employed in these studies (CK+DCI) has been optimized to support iAT2 differentiation and proliferation, and thus may have increased this iAT1-iAT2 reversion; however, overall, our findings suggest the necessity for activated nuclear YAP in the maintenance of AT1 program in human systems as well.

Our results raise several questions that will require further study. First, while our defined medium leads to robust differentiation of iAT2s into iAT1s by single cell RNA sequencing, it remains unclear whether this transition proceeds through the recently described transitional or basaloid cell state.^2, 3, 66, 67^ Although a very minor subset of iAT1s at the end point of our experiments expressed a few markers potentially associated with the transitional state (e.g. *KRT17*), most transitional markers were absent and further kinetic studies are needed at single cell resolution, potentially including bar-coded lineage tracing approaches, to understand whether differentiation fate trajectories in our model include transitional or other intermediate phenotypes. A second priority in future work will be further maturing the iAT1s produced by our methods. While our iAT1s show some functional properties and transcriptomic similarities to that of in vivo human AT1 cells, expression levels of most canonical AT1 markers are still significantly lower than in primary cell controls, and a variety of future approaches can now be tested in an effort to augment maturation, such as introducing biomechanical cues or different matrix substrata that more closely recapitulate the distal lung microenvironment. Lastly, our reductionist model which includes only 2 cell types in our studies (iAT1s and/or iAT2s), can be augmented in complexity in the future, beginning with the introduction of other alveolar lineages into co-cultures that might include mesenchymal, vascular, and immune lineages.

Thus, our work shows the generation of AT1-like cells from iPSC-derived AT2 cells, providing an in vitro model of human alveolar epithelial differentiation and a potential source of human cells that until now have been challenging to viably obtain from patients. Access to these cells, either in pure form or combined with other lineages should facilitate a variety of basic developmental studies, disease modeling, and potential engineering of future regenerative therapies.

## Methods

### iPSC generation and maintenance

The reprogramming and characterization of the original human iPSC clones employed in this study (BU3 and SPC2-ST-B2; aka “SPC2B2”) were previously published.^55, 59, 64^ All iPSC lines had a normal karyotype (Cell Line Genetics G-banding analysis), both before and after gene-editing, and were maintained on hESC qualified Matrigel (Corning, 8774552) in feeder-free conditions in mTeSR1 medium (STEMCELL Technologies, 05850). Gentle Cell Dissociation Reagent (STEMCELL Technologies, 07174) was used for passaging. All iPSC differentiations were performed under regulatory approval of the Institutional Review Board of Boston University. Additional details and protocols for iPSC derivation, culture, and characterization can be downloaded at https://crem.bu.edu/cores-protocols/. All iPSC lines are available from the CReM repository upon request, https://stemcellbank.bu.edu.

### CRISPR targeting of tdTomato to AGER locus in iPSCs

The iPSC line BU3 NG,^59^ previously engineered to carry an NKX2-1^GFP^ reporter was used for targeting a tdTomato reporter to the human AGER endogenous locus using CRISPR gene editing as follows. Left and right homology arms of about 700bp in length to the left and right of the AGER endogenous start codon were generated by PCR amplification using gDNA extracts from BU3 NG iPSCs. These arms were cloned into our previously published plasmid backbone p1303-DV- SFTPC-tdTomato^29^ replacing the SFTPC homology arms, generating the p2701-AGER-tdTomato plasmid. Guide RNAs (gRNA1: CACCGCCAGGCTCCAACTGCTGTTC; gRNA2: CACCGATGGCTGCCGGAACAGCAGT; gRNA3: CACCGCTGTGGCCTCCGCCCTAGGT) were selected and cloned into the pSpCas9(BB)-2A-GFP plasmid (Addgene plasmid #48138). BU3 NG iPSCs were pretreated with ROCK Inhibitor (Y-27632) for three hours prior to nucleofection using the Lonza P3 Primary Cell 4DNucleofector™ X Kit (Lonza, cat. no. V4XP-3024) and replated for puromycin resistance screening. Clones were passaged and gDNA was isolated for PCR screening using the following primer pairs (Fig S5): 5’ CTGATCCCCTCAGACATTCTCAGGA 3’ to 5’ GAGCTGCCGCTGCCGGT 3’ for outside the left homology arm to the tdTomato cassette and 5’ ACTTGTGTAGCGCCAAGTGC 3’ to 5’ ACACACACTCGCCTCCTGTT 3’ for within the puromycin resistance cassette to outside the right homology arm. HEK293 cells that had been transfected with the targeting plasmids or not were used as controls. Clones that showed insertion of tdTomato by PCR were sequenced to confirm insertion, and further expanded. The floxed PGK-Puromycin resistance cassette was excised using transient transfection with Cre plasmid (pHAGE2 EF1aL-Cre-IRES-NeoR-W; plasmid map available at www.kottonlab.com) as previously published^29^ with transient G418 selection of candidate targeted, Cre-excised clones. Mono-allelic targeting and puromycin resistance excision was confirmed by PCR with primers 5’AGGACTCTTGTCCCAAAGGC 3’ to 5’ CTGGGGTGTGGGGTTAAAGT 3’ yielding both a 267bp long band for the untargeted allele and a 2290bp long band for the targeted allele with the puromycin cassette excised. Cells were then assessed by G-banding to identify karyotypically normal clones (Cell Line Genetics).

### iPSC differentiation into iAT2 cells (Alveolospheres)

Human iPSC lines (BU3 NG, BU3 NGAT, SPC2-ST-B2 were differentiated into iAT2s as previously described^69^, with detailed protocols and characterizations available for free download at www.kottonlab.com. Briefly, the STEMdiff Definitive Endoderm Kit (STEMCELL Technologies, 05110), was used for differentiation into endoderm, which was scored by co-expression of CKIT and CXCR4 by flow cytometry. Cells were then passaged using Gentle Cell Dissociation Reagent, replated onto Matrigel coated plate, and cultured in “DS/SB” media consisting of complete serum free differentiation media (cSFDM)^69^ with 2uM Dorsomorphin (“DS”; Stemgent) and 10uM SB431542 (“SB”; Tocris) for 72 hours for anteriorization, the first 24 hours being supplemented with 10 uM Y-27632 (Tocris). Media was then changed to cSFDM with 3 uM CHIR99021 (“C”; Tocris), recombinant human BMP4 (10 ng/ml; “B”; R&D Systems), and 100 nM retinoic acid (“Ra”; Sigma-Aldrich) called “CBRa” for lung specification. Between days 14-16 of differentiation, cells were sorted for lung progenitors either using NKX2-1^GFP^ knock-in reporters or using antibody staining for CPM (FUJIFILM) for lines not containing an NKX2-1 reporter. Sorted cells were resuspended in growth-factor reduced Matrigel (Corning 356231) droplets and covered with alveolar differentiation medium, “CK+DCI” containing a base of cSFDM with 3 uM CHIR99021 (C), rhKGF (10 ng/m; “K”; R&D Systems), 50 nM dexamethasone (“D”; Sigma-Aldrich), 0.1 mM 8-bromoadenosine 3′,5′ cyclic monophosphate sodium salt (Sigma-Aldrich), and 0.1 mM 3- isobutyl-1-methylxanthine (IBMX; Sigma-Aldrich) (“CI”). 10 uM Y-27632 (Tocris) was supplemented for 72 hours post sort and cells were refed with CK+DCI every 48-72 hours. For iAT2 maintenance, cells were passaged every 10-14 days as single cells as previously described.^69^ iAT2s at Air-liquid interface were plated onto 6.5mm transwell inserts (Corning) coated with hESC qualified Matrigel (Corning, 8774552) as previously published.^68, 70^

### Lentiviral and Adenoviral Transduction

For introduction of lentiviral and adenoviral constructs to iAT2s, alveolospheres were dissociated to single cells as with passaging^69^. Cells were then incubated for 4 hours at 37C in suspension with virus in CK+DCI supplemented with 10 uM Y-27632 and 5ug/mL polybrene. WT YAP and YAP5SA lentiviral transduction was performed at a multiplicity of infection (MOI) of 10, as previously published.^69^ An MOI of 200 was used for infections with Adeno-Cre-GFP virus. Cells were then replated in Matrigel droplets in CK+DCI.

### iAT1 differentiation in “L+DCI” medium and air-liquid interface (ALI) culture

iPSCs were first differentiated into iAT2s and passaged as above in 3D cultures in iAT2 medium (CK+DCI) supplemented for the first 3 days after passaging with 10 uM Y-27632 (Tocris). To generate iAT1s, 3 days after passaging iAT2s in 3D, the medium was replaced with “L+DCI”, consisting of [10 uM LATS-IN-1 (“L”; MedChemExpress Cat. No.: HY-138489), 50 nM dexamethasone (“D”; Sigma-Aldrich), 0.1 mM 8-bromoadenosine 3′,5′ cyclic monophosphate sodium salt (Sigma-Aldrich; “C”), and 0.1 mM 3-isobutyl-1-methylxanthine (IBMX; Sigma-Aldrich; “I”)]. Cells were cultured up to 16 days in L+DCI in 3D while monitoring AGER^tdTomato^ expression as detailed in the text.

ALI versions of iAT1 cultures were prepared as follows. To prepare “iAT1 ALI p0” cultures, first a single cell suspension of iAT2s was passaged onto 6.5mm transwell inserts (Corning) coated with hESC qualified Matrigel (Corning, 8774552) as previously published,^68, 70^ but switching CK+DCI medium to L+DCI at the time of plating. To prepare “iAT1 ALI p1” cultures, iAT2s in 3D Matrigel culture were first switched to L+DCI medium in 3D for 9 days before being passaged as single cells onto Matrigel coated transwells for continued L+DCI culture. (High density = 200k cells/6.5mm insert, medium = 100k cells/insert, low density = 50k cells/insert.) For all ALI culturing conditions, rock inhibitor was added to LDCI media for the first 3 days post passaging and then removed. At the same time (day 3), liquid was aspirated from the apical chamber (air lift) to form the air liquid interface (ALI). Cells in ALI cultures were maintained for up to 10 days (7 days post air lift). Accutase (Innovative Cell Technologies) was used to dissociate cells for FACS analysis and single cell RNA sequencing.

### Primary AT2 culture and differentiation

Primary AT2 cells, a generous gift of Purushothama Rao Tata (Duke University), were cultured in Serum Free Feeder Free (SFFF) medium in 3D Matrigel as reported.^32^ For differentiation experiments, cells were passaged as reported and cultured in SFFF for 7 days before media was switched to either L+DCI or published human serum containing Alveolar Differentiation Medium (ADM).^32^

### Reverse Transcription quantitative polymerase chain reaction (RT-qPCR)

For RT-qPCR, whole well or sorted cells were collected and stored in Qiazol (Qiagen, 79306) prior to RNA isolation using RNeasy Plus Mini Kit according to the manufacturer’s protocol (Qiagen, 74104). cDNA was then synthesized using MultiScribe™ Reverse Transcriptase (ThermoFisher 4311235). A QuantStudio instrument (Applied Biosciences) and predesigned Taqman probes were used and run in a 384-well format for 40 cycles. Relative expression was normalized to an 18S control and fold change over control cells was calculated using 2^-ΔΔCt^. Where indicated in the text, RNA extracts from adult primary human distal lung tissue explants, the kind gift of Barry Stripp (Cedars Sinai, Los Angeles) were employed as RT-qPCR controls. RNA was isolated via RNeasy Plus Mini Kit following manufacturer’s protocol.

### Flow Cytometry and FACS

0.05% trypsin was used to generate single cell suspensions which were resuspended in sort buffer [HBSS (ThermoFisher) with 2% FBS, 10 uM Y-27632 (Tocris)] with Live/dead stain [Calcein blue (Life technologies) or DRAQ7 (Abcam)]. Cells were sorted based on reporter expression: NKX2-1^GFP^ and AGER^tdTomato^ for BU3 NGAT, or SFTPC^tdTomato^ for SPC2B2, as indicated in the text. Cell sorting was performed on a Moflo Astrios EQ (Beckman Coulter) and flow cytometry analysis was performed on an LSRII SORP (BD Biosciences) at the Boston University Flow Cytometry Core Facility. For Edu assays, the Click-iT Plus EdU Alexa Fluor 647 Flow Cytometry Assay Kit (Thermo Fisher Scientific) was used with EdU added 24 hours before cell isolation, and cells fixed in 4% paraformaldehyde (PFA) were analyzed on a Stratedigm (S1000EXI) cytometer with post-processing using FlowJo software (BD Biosciences). For all other analyses, live non-fixed cells were sorted or analyzed as indicated in the text.

### Immunofluorescence microscopy

Organoids (3D cultured epithelial spheres or clumps) or cells were fixed in 4% PFA for 20 minutes at room temperature. For whole mount staining, organoids were immediately stained in solution and mounted on cavity slides (Elisco). For sectioning, samples were dehydrated via ethanol and paraffin embedded, or dehydrated using sucrose and frozen in OCT. Prior to staining, paraffin sections were rehydrated, and antigen retrieval was performed using heated citrate buffer. For both paraffin- and cryo-embedded sections, permeabilization was performed using 0.1% Triton and blocking was done with 4% Normal Donkey Serum. Samples were incubated in primary antibody overnight at 4C and incubated in secondary antibody with Hoechst for 2 hrs at RT. Primary antibodies used included ProSFTPC (Pro-SPC): Seven Hills WRAB-9337; HTI-56: Terrace Biotech TB-29AHT1-56; ZO-1: Thermo Fisher Cat# 61-7300; and RFP: Rockland 600- 401-379. Confocal images were taken on either a Leica SP5 or Zeiss LSM 710-Live Duo and were processed using Fiji.

For ALI cross sections, cultures were fixed for 2 hours at RT in 2% glutaraldehyde plus 1% paraformaldehyde in 0.1M Cacodylate buffer, pH 7.4 then post fixed overnight at 4C in 1.5% osmium tetroxide. After washing, membranes were cut away from the insert, placed in glass vials and block stained in 1.5% uranyl acetate. Samples were dehydrated through graded acetones, infiltrated in a 50:50 mixture of propylene oxide: resin and embedded in Embed 812. 0.5-1um sections were cut and stained with Toluidine blue.

### VEGFA ELISA

To measure secreted VEGFA protein, conditioned media after 48 hours of exposure to cells was harvested from each sample indicated in the text and figure legends on day 10 of ALI culture. VEGFA ELISA was performed using the Human VEGFA ELISA kit (Abcam) according to manufacturer’s instructions.

### Cell surface area calculations

iAT1 ALI p1 cells were grown in LDCI medium for 10 days, aspirating apical medium on day 3 (air lift) to form an air liquid interface. Day 10 post plating (7 days after air lift) cells were fixed in 4% PFA and stained for tight junction protein ZO-1 (Thermo Fisher Cat# 61-7300). Randomized Z stack images were taken at 20x in 3 different places of each transwell for three transwells and imported into ImageJ. Cell outlines denoted by ZO-1 were traced by hand and area was calculated using ImageJ for around 50 cells per image.

#### Transepithelial Electrical Resistance

To measure TEER, a Millicell ERS-2 Coltohmmeter (Moillipore Sigma, MERS00002) was used. Electrodes were sterilized by dipping in 70% EtOH followed by conditioned cell media. 200uL media was added to the apical chamber of Transwell culture inserts prior to taking measurements. For each sample, readings were taken at 3 locations in the well. TEER was calculated by subtracting the “blank” Matrigel coated well from the mean and multiplying by the tissue culture growth area.

#### Generation of YAP5SA and WT YAP lentivirus

Coding sequences for YAP5SA and WT YAP were PCR amplified from pLVX-Tight-Puro-3F-YAP- 5SA and pLVX-Tight-Puro-3F-YAP, respectively,^71^ adding NotI and BglII restriction sites for ligation into a pHAGE2-EF1aL-dsRed-UBC-tagBFP-WPRE backbone in place of the dsRed cassette. We have previously published the dual promoter lentiviral vector for dual transgenesis^72^ and the pHAGE2 version includes a loxP cassette in the 3’LTR as previously published.^73^ This cloning strategy generated the following loxP containing plasmids for the current project: pHAGE2-EF1aL-YAP5SA-UBC-tagBFP-WPRE and pHAGE2-EF1aL-WTYAP-UBC-tagBFP-WPRE. The YAP5SA coding sequence was also cloned into a pHAGE1 backbone generating pHAGE1-EF1aL-YAP5SA-UBC-GFP-WPRE. Lentiviral packaging and titering protocols (as we have previously published),^72^ as well as plasmid maps and primer design protocols for cloning into pHAGE lentiviral backbones are all available at www.kottonlab.com.

#### Single Cell RNA sequencing of iAT2s and iAT1s

For single cell RNA sequencing of iPSC-derived alveolar epithelial cells, cells were dissociated to single cell suspensions using 0.05% trypsin. Live cells were sorted via DRAQ7 using Moflo Astrios EQ (Beckman Coulter) at the Boston University Flow Cytometry Core Facility. Single Cell RNA sequencing was then preformed using the Chromium Single Cell 3’system according to manufacturers’ instructions (10x Genomics) at the Single Cell Sequencing Core at Boston University Medical Center. Sequencing files were mapped to the human genome reference (GRCh37) supplemented with GFP, tdTomato, and BFP sequences using CellRanger v3.0.2. Seurat v3.2.3^74^ was used for downstream analysis and quality control. After inspection of the quality control metrics, cells with 15% to 35% of mitochondrial content and <800 detected genes were excluded for downstream analyses. In addition, doublets were also excluded for downstream analysis. We normalized and scaled the unique molecular identifier (UMI) counts using the regularized negative binomial regression (SCTransform).^75^ Following the standard procedure in Seurat’s pipeline, we performed linear dimensionality reduction (principal component analysis) and used the top 20 principal components to compute the unsupervised Uniform Manifold Approximation and Projection (UMAP).^76^ For clustering of the cells, we used Louvain algorithm^77^ which were computed at a range of resolutions from 1.5 to 0.05 (more to fewer clusters). Populations were annotated using Louvain Clustering resolution indicated in the text. Cell cycle scores and classifications were done using the Seurat’s cell-cycle scoring and regression method.^78^ Cluster specific genes were calculated using MAST framework in Seurat wrapper.^79^ An online Shiny app^80^ has been established to allow interactive, user-friendly visualizations of gene expression in each population that will be available on the Kotton Lab’s bioinformatics portal www.kottonlab.com.

#### Primary Single Cell RNA sequencing

##### Tissue preparation and scRNA sequencing

Samples of normal de-identified human lungs from donors who were not matched for lung transplant, were obtained as described previously^41^ with the following adaptations. For peripheral tissue experiments, a 3 × 2 cm piece of distal lung tissue was obtained, pleura and visible airways/blood vessels were dissected away, mechanically minced into ∼2mm pieces, and processed into a single-cell suspension as described.^41^ After a single-cell suspension was obtained from the proximal or peripheral tissue, CD45+ immune cells were depleted via MACS LS columns using CD45-microbeads (Miltenyi, 130-045-801) with 2 X 10^6^ cells per column to enhance purity and viability. After CD45 depletion, sorted cells were loaded onto a GemCode instrument (10x Genomics, Pleasanton, CA, USA) to generate single-cell barcoded droplets (GEMs) according to the manufacture’s protocol. The resulting libraries were sequenced on an Illumina HiSeq2500 or NovaSeq instrument.

##### Analysis of scRNA-seq data

Read were aligned and unique molecular identifier (UMI) counts obtained via the Cell Ranger pipeline (10X Genomics). For further processing, integration and downstream analysis, Seurat V3 (PMID: 29608179) was used. Cells with less than 200 genes, greater than 2 Median absolute deviation above the median, and with potential stress signals of greater than 10% mitochondrial reads were removed. The Cell Cycle phase prediction score was calculated using Seurat function CellCycleScoring, and data was normalized and scaled using the SCTransform function and adjusting for cell cycle, percent mitochondria, number of features per cell, and number of UMI per cell. For integration of individual samples, Canonical Correlation Analysis (CCA) method using SCTransform in Seurat V3; the top 3000 variable genes were used to find anchors. Linear dimension reduction was done via PCA, and the number of PCA dimensions was evaluated and selected based on assessment of an ElbowPlot. Data was clustered using the Louvain graph based algorithm in R and cluster resolution was evaluated using R package ‘clustree’. The Uniform Manifold Projection (UMAP) data reduction algorithm was used to project the cells onto two dimensional coordinates. Subsequently canonical marker genes were used to identify cellular compartments (epithelium, endothelium, mesenchymal, and immune populations). Epithelial clusters were subsetted, and clustering and UMAP reduction were repeated. Clusters were then assigned putative cell types based on annotation with canonical marker genes, or from assessment of top cluster-defining genes based on differential expression (using the FindConservedMarkers function in SueratV3).

#### Development of primary human AT1 gene signatures

Differential gene expression analysis was performed using the Seurat function FindAllMarkers using the MAST algorithm and using the RNA assay for UMIs. The following comparison were performed 1) AT1s vs all lung cells, 2) AT1s vs. lung epithelial cells, and 3) AT1s vs AT2s. The top 50 differentially upregulated genes ordered by LogFC (FDR<0.05) were used as AT1 gene signatures (Table S1), with AT1 vs all lung cells being used as the AT1 gene signature scored in Figures 3, 6, and 7. Similarly, an AT2 50-gene signature (vs all lung cells) was generated and scored in figure 6. AT1 gene signature (and other signatures) are made using module score function of Seurat

### Statistical analysis

Unpaired student’s t-tests were used to compare two groups while one-way Analysis of Variance (ANOVA) with Tukey multiple comparisons test was used to compare the means between three or more groups. p<0.05 was used to determine statistical significance unless otherwise indicated in the text.

### Data Availability

Primary scRNA-seq data analyzed during this study have been deposited at the Gene Expression Omnibus (GEO) database (accession numbers <u>GSE168191</u>) iPSC-derived alveolar cell scRNA seq datasets have been deposited at the GEO database with the following accession numbers: Fig 3 WT YAP,YAP5SA: GSE221342; Fig 6 LDCI, YAP5SA, CKDCI: GSE221343; Fig 7 3D vs ALI: GSE221344

### Study Approval

All iPSC differentiations were performed under regulatory approval of the Institutional Review Board of Boston University (protocol H33122).

## Author Contributions

CLB and DNK conceived the project and designed experiments. EM and XV provided expert input on experimental design and analysis. CLB, JH, KDA, BRT, KM, AH, and AMM provided resources and performed experiments. PB, MM, AB, CVM, and FW analyzed sequencing data. CLB and DNK wrote the manuscript. All authors reviewed and approved final version.

## Acknowledgements

The authors wish to thank all members of the Kotton, Varelas and Morrisey Labs for insightful discussions. We thank Yuriy Alekseyev of the Boston University School of Medicine (BUSM) Single Cell Sequencing Core, supported by NIH grant 1UL1TR001430, and Brian R. Tilton of the BUSM Flow Cytometry Core. We are grateful to Greg Miller and Marianne James of the Boston University Center for Regenerative Medicine (CReM) for maintenance and characterization of patient-specific iPSCs, supported by NIH grants NO1 75N92020C00005 and U01TR001810. We thank Purushothama Rao Tata (Duke University, Durham, NC) for the generous gift of cultured human primary AT2 cells. The schematics were created with BioRender.com. This study was supported by National Institutes of Health grants 1F31HL158193, T32HL007035, and a PCTC JumpStart Award PCTC_JC_202003 to C.B.; an I.M. Rosenzweig Junior Investigator Award from The Pulmonary Fibrosis Foundation and an Integrated Pilot Grant Award through Boston University Clinical & Translational Science Institute (1UL1TR001430) to K.D.A.; R01HL124392 to X.V.; U01HL134745, U01HL134766, U01HL152976, and R01HL095993 to D.N.K. iPSC derivation and cell sharing was supported by N01 75N92020C00005 and U01TR001810 to D.N.K.

---

**Supplemental Figure 1:**
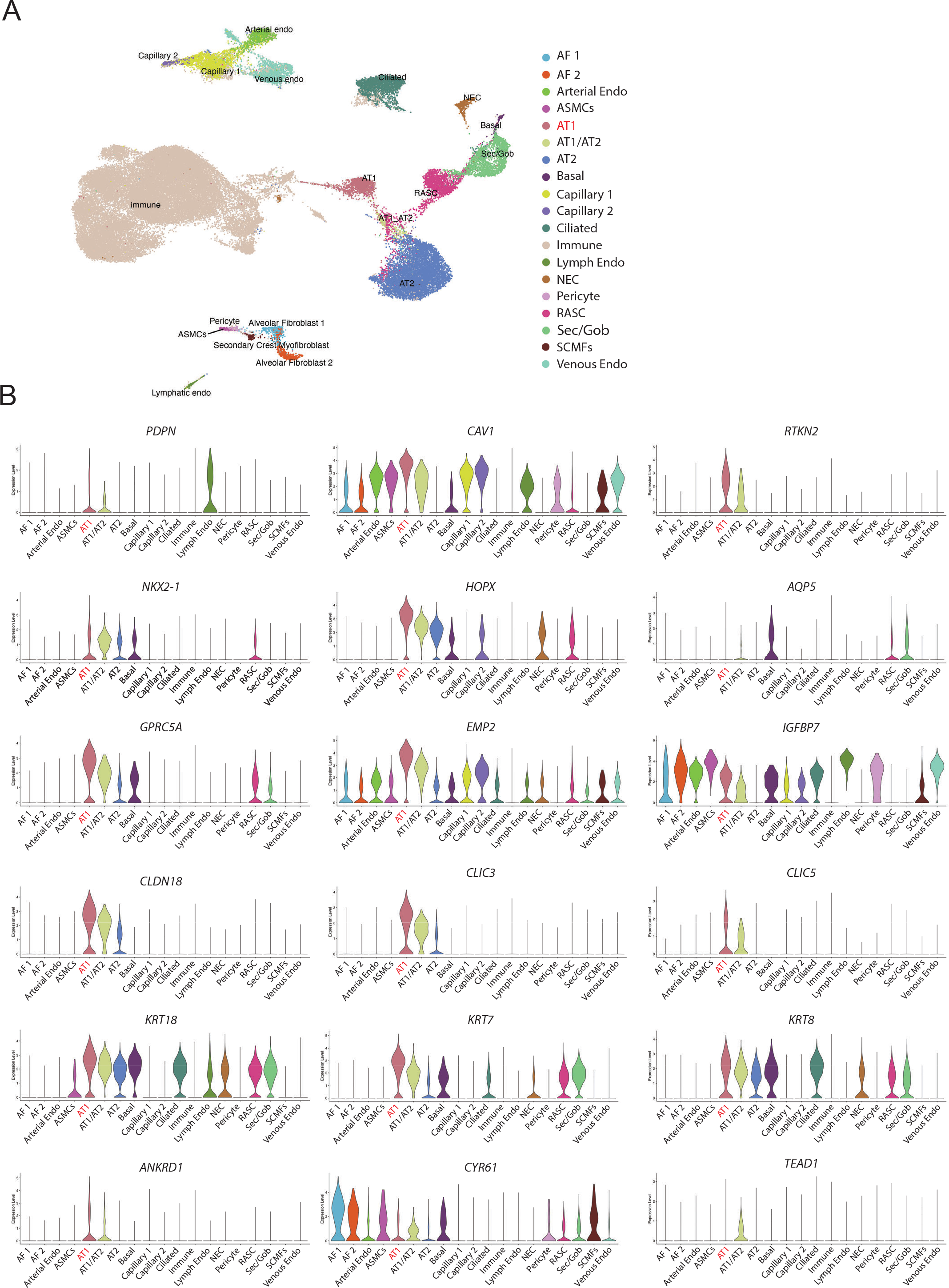
Human AT1 marker gene expression across all cell types in the lung. A) UMAP projection of all lung cells in previously published dataset by Basil et al.^41^ (N=5, 58567 total cells, 1401 AT1s). B) Violin plots showing gene expression of selected AT1 marker genes across all lung cell types represented in the dataset.

**Supplemental Figure 2:**
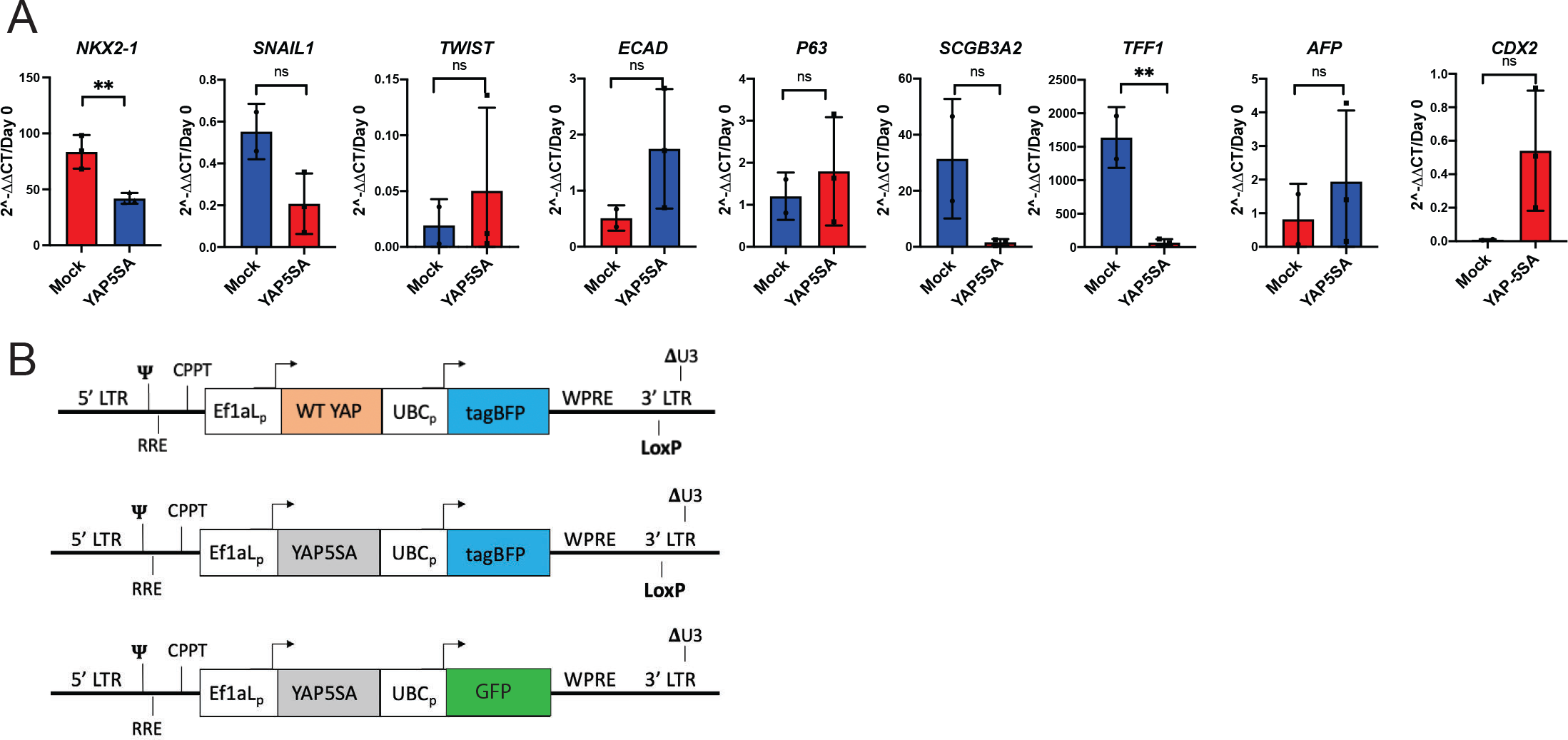
YAP5SA transduction of SPC2B2 iAT2s. A) Whole well gene expression by RT-qPCR of EMT markers, airway lung lineage markers and non-lung endoderm markers 14 days post YAP5SA or mock lentiviral transduction of SPC2B2 iAT2s, relative to Day 0 iPSCs. (N=3 per condition, student’s t test) B) Diagram of WT YAP and YAP5SA lentiviruses showing dual promoter system with tagBFP and LoxP site. YAP5SA-GFP lentivirus used for competition assay. *p<0.05, **p<0.01, ***p<0.001, and ****p<0.001 for all panels.

**Supplemental Figure 3:**
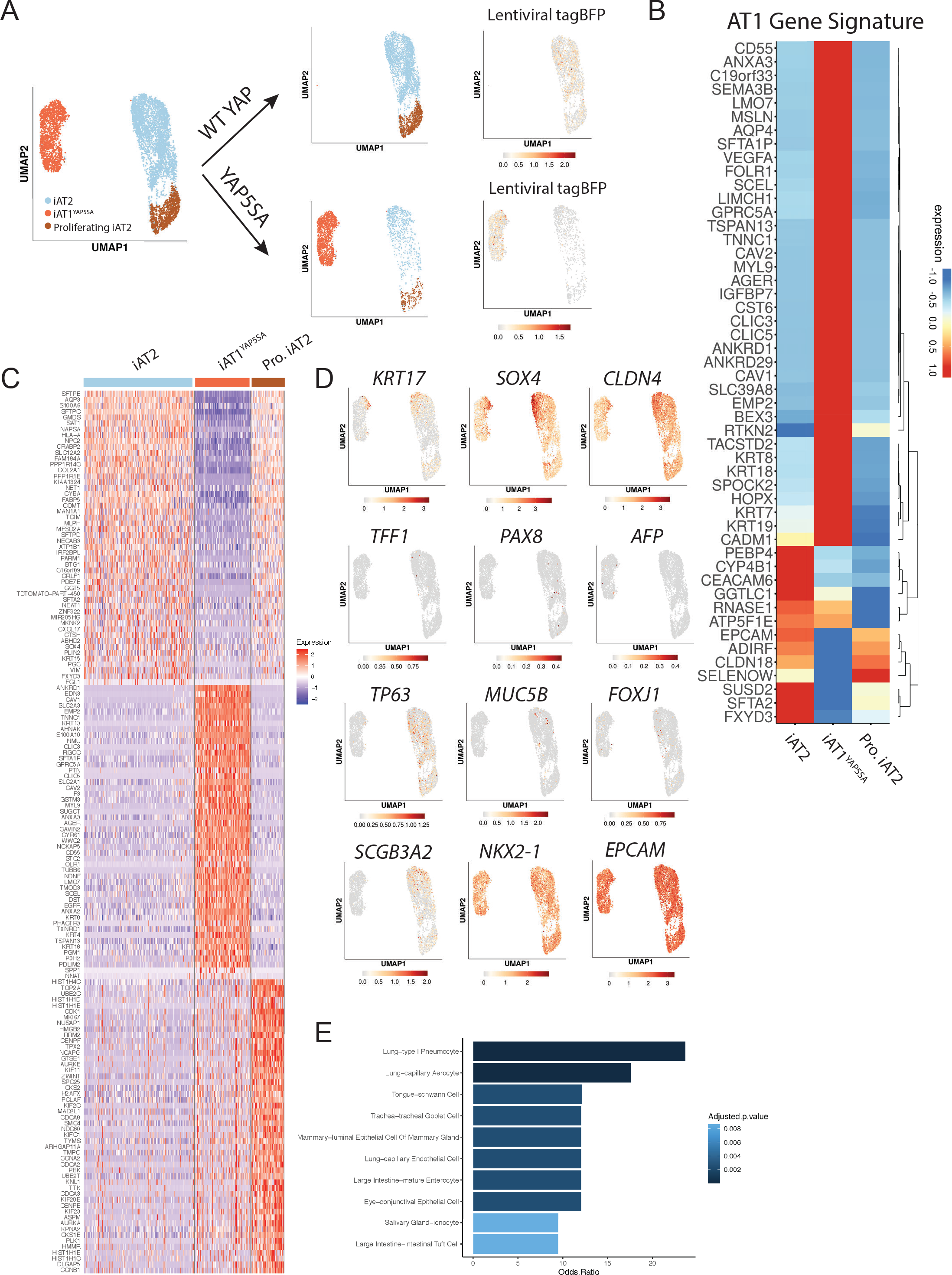
Single Cell RNA sequencing of YAP5SA transduced iAT2s. A) Sub plots of UMAP projections of WT YAP and YAP5SA wells to show lentiviral tagBFP expression in each condition. B) Heatmap of genes in the 50 AT1 gene signature across each population. C) Full heatmap of top 50 differentially upregulated genes in the iAT2, iAT1^YAP5SA^, and Proliferating iAT2 populations. D) Gene expression overlays of aberrant basaloid transitional markers *KRT17, SOX4*, and *CLDN4*; Non-lung endoderm markers *TFF1, PAX8*, and *AFP*; and other lung markers *TP63, MCU5B, FOXJ1, SCGB3A2*, and *NKX2-1*. E) Enrichr analysis of top 92 (FDR > 0.05, logFC > 1) differentially upregulated genes in iAT1^YAP5SA^ population using Tabula Sapiens dataset.

**Supplemental Figure 4:**
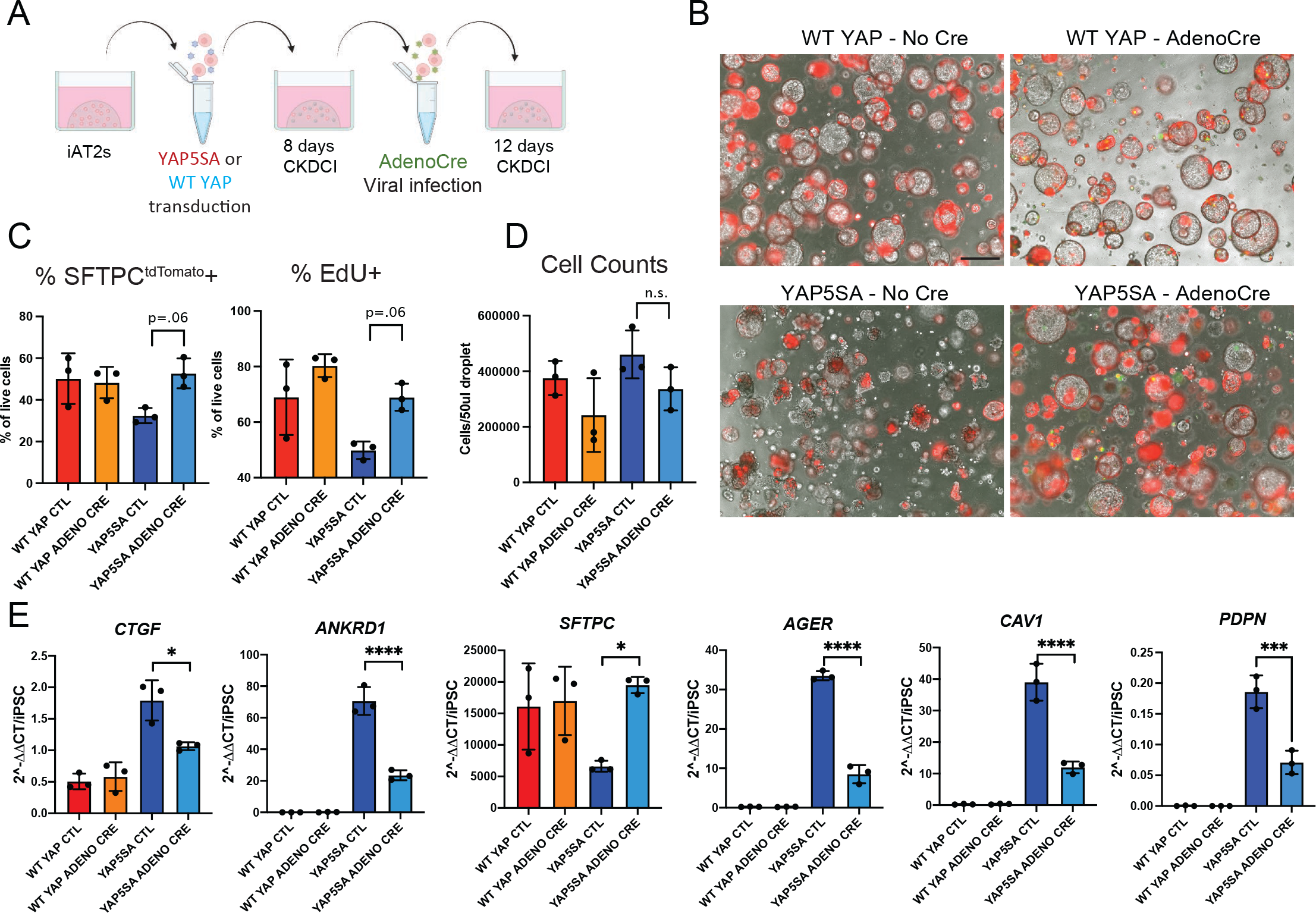
Cre excision of YAP5SA lentivirus leads to reversion to iAT2-like phenotype. A) iAT2s were transduced with either WT YAP or YAP5SA lentivirus and then plated in 3D Matrigel in CK+DCI for 8 days. They were then infected in single cell suspensions with either Adeno-Cre to excise lentivirus or mock and replated into 3D Matrigel in CK+DCI for 12 days. B) Representative live cell imaging of SPC2-ST-B2 iAT2s following AdenoCre infection (SFTPC^tdTomato^/Phase Contrast overlay, scale bar = 500um). C) Quantification of flow cytometry analysis showing proliferation by 24hr EDU and SFTPC^tdTomato^ percentage. (N=3, 1-way ANOVA). D) Cell counts per 50uL droplet. E) Gene expression of YAP downstream targets, AT2 and AT1 markers by bulk RT-qPCR (N=3, 1-way ANOVA). *p<0.05, **p<0.01, ***p<0.001, and ****p<0.001 for all panels.

**Supplemental Figure 5:**
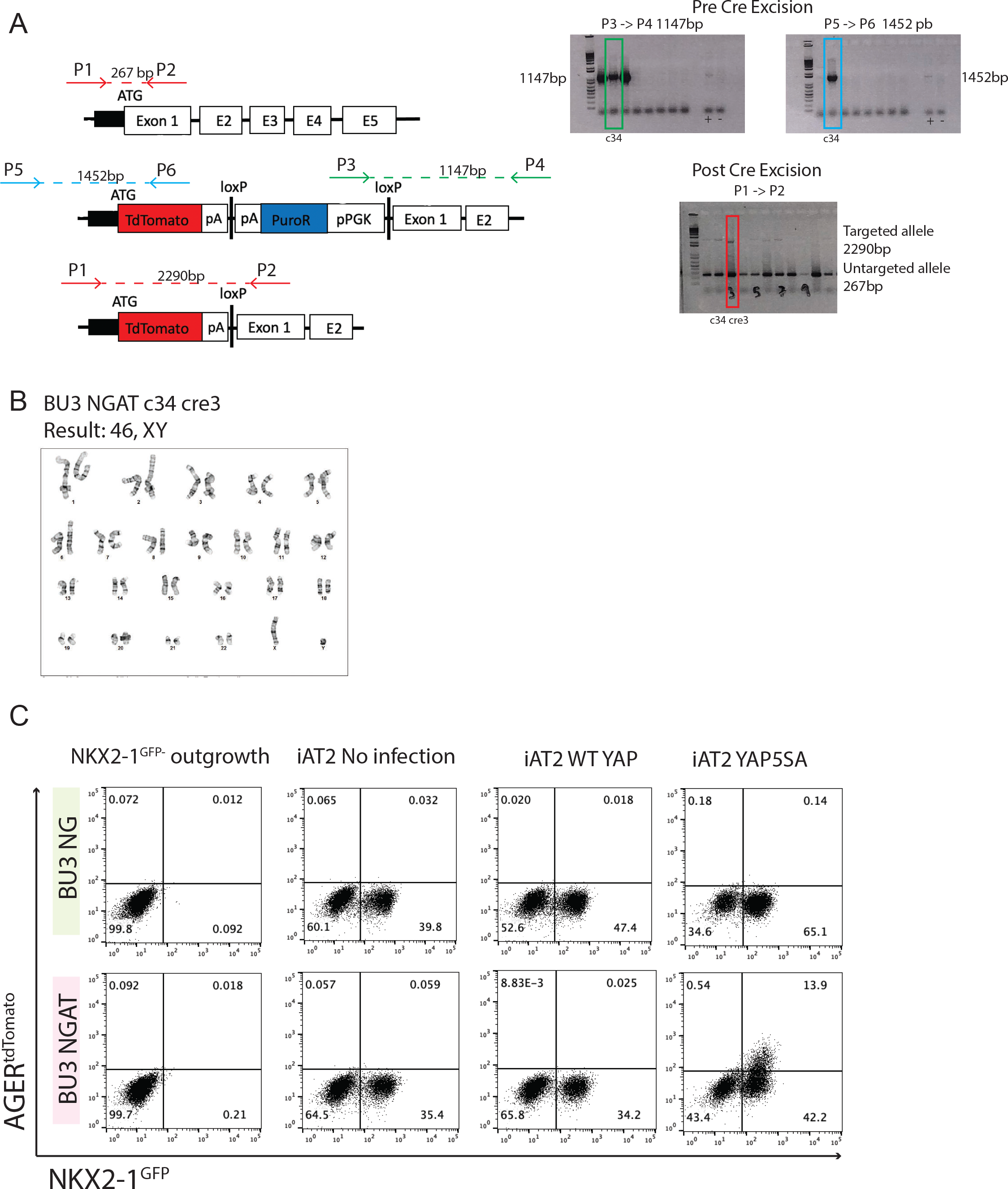
Gene editing and characterization of a NKX2-1^GFP^/AGER^tdTomato^ dual reporter iPSC line. A) CRISPR targeting of tdTomato reporter to *AGER* locus and Cre excision of puromycin resistance cassette. Final clone does not include puromycin cassette by lack of Fp2 -> Rp2 band and has both long and short bands from Fp1 -> Fp2 showing one copy of unedited *AGER* and one copy of tdTomato reporter. B) Karyotype of BU3 NGAT iPSC line. C) Representative flow cytometry of NKX2-1^GFP^ negative outgrowth, iAT2s, WT YAP transduced and YAP5SA transduced cells in both BU3 NGAT and parent BU3 NG lines.

**Supplemental Figure 6:**
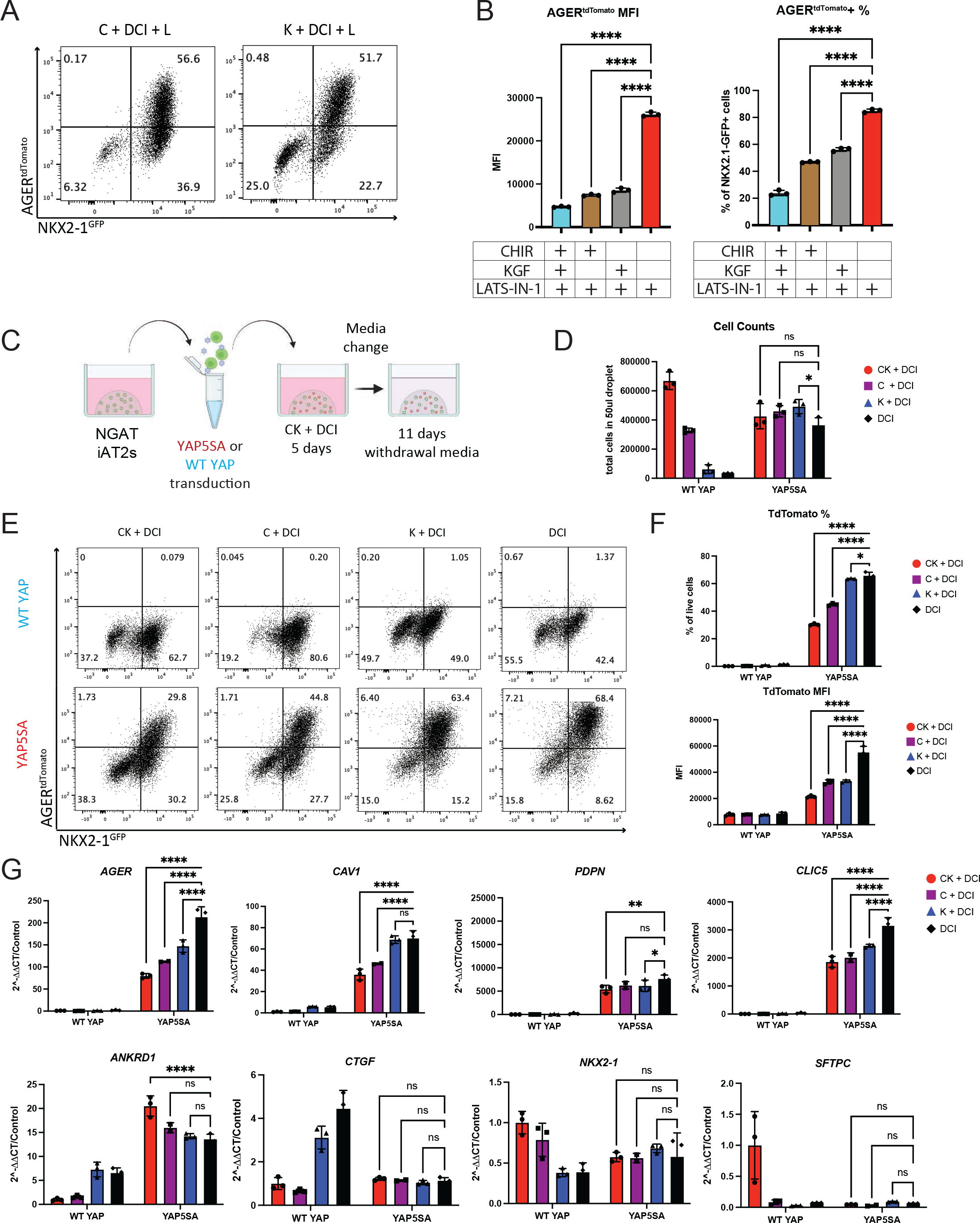
Both Chir and KGF are inhibitory towards AT1 program induced by both serum-free media and YAP5SA lentivirus. A) BU3 NGAT iAT2s were passaged into 3D Matrigel in CK+DCI for 3 days, and then media was switched to media containing LATS-IN-1 and either Chir or KGF and grown for a further 11 days. Representative flow cytometry of NKX2-1^GFP^ and AGER^tdTomato^. B) Quantification of AGER^tdTomato^ compared to CK DCI + L and DCI + L quantification in Figure 5. (N=3 per condition, 1-way ANOVA). C) BU3 NGAT iAT2s were transduced with either WT YAP or YAP5SA lentivirus in suspension for 4 hours and then replated into 3D Matrigel in CK DCI. After 5 days, media was changed to withdraw one or both growth factors from the media. Cells were analyzed 11 days post media change. D) Cell counts of WT YAP and YAP5SA transduced cells in different medias (2-way ANOVA). E) Representative flow cytometry of NKX2-1^GFP^ and AGER^tdTomato^. F) Quantification of AGER^tdTomato^ percentage and Mean Fluorescence Intensity following YAP5SA transduction. (2-way ANOVA). G) Gene expression of AT1 markers, YAP downstream targets, and AT2 markers by whole well RT-qPCR. (N=3, 2-way ANOVA). p<0.05, **p<0.01, ***p<0.001, and ****p<0.001 for all panels.

**Supplemental Figure 7:**
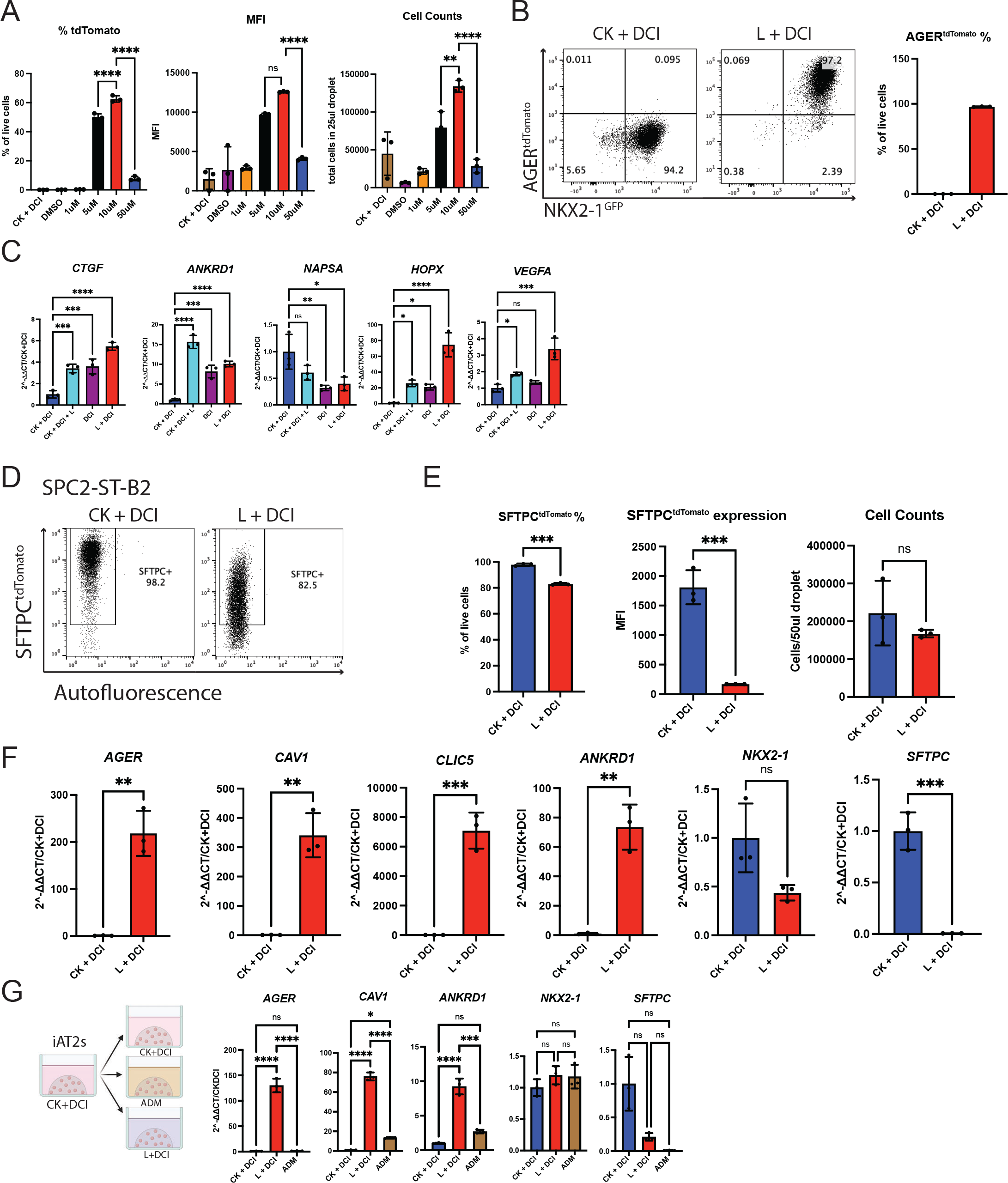
Optimization of LATS inhibitor-containing media for iAT1 induction. A) Quantification of AGER^tdTomato^ expression by flow cytometry and cell counts of different concentrations of LATS-IN-1 added 3 days post passage, analyzed 9 days post media change. (N=3 per condition, 1-way ANOVA). B) Flow cytometry analysis and quantification of AGER^tdTomato^ in BU3 NGAT cells that were sorted on NKX2-1^GFP^ on day 201 and replated into CK+DCI+RI before media was switched to L+DCI 3 days later. Cells were analyzed 11 days post media change. (N=3). C) Gene expression of YAP downstream targets, and AT2 and AT1 markers in LATS inhibitor media by whole well RT-qPCR. (1-way ANOVA). D) SPC2-ST-B2 iAT2s (>95% SFTPC^tdTomato^+) were passaged into CK+DCI and then medium was switched to L+DCI 3 days post passage. Representative flow cytometry of SFTPC^tdTomato^ reporter expression 8 days post media change. E) Quantification of SFTPC^tdTomato^ and cell counts of SPC2-ST-B2 cells in CK+DCI and L+DCI (N=3 per condition, students t test). F) Gene expression by bulk RT-qPCR at 8 days post media change. (N=3) G) BU3 NGAT iAT2s were cultured in CK+DCI for three days post passage before changing to Alveolar Differentiation Medium (“ADM”; Katsura et al. 2020)^32^ or L+DCI and whole well RNA was taken for RT-qPCR 7 days post media change. (N=3, One-way ANOVA). p<0.05, **p<0.01, ***p<0.001, and ****p<0.001 for all panels.

**Supplemental Figure 8:**
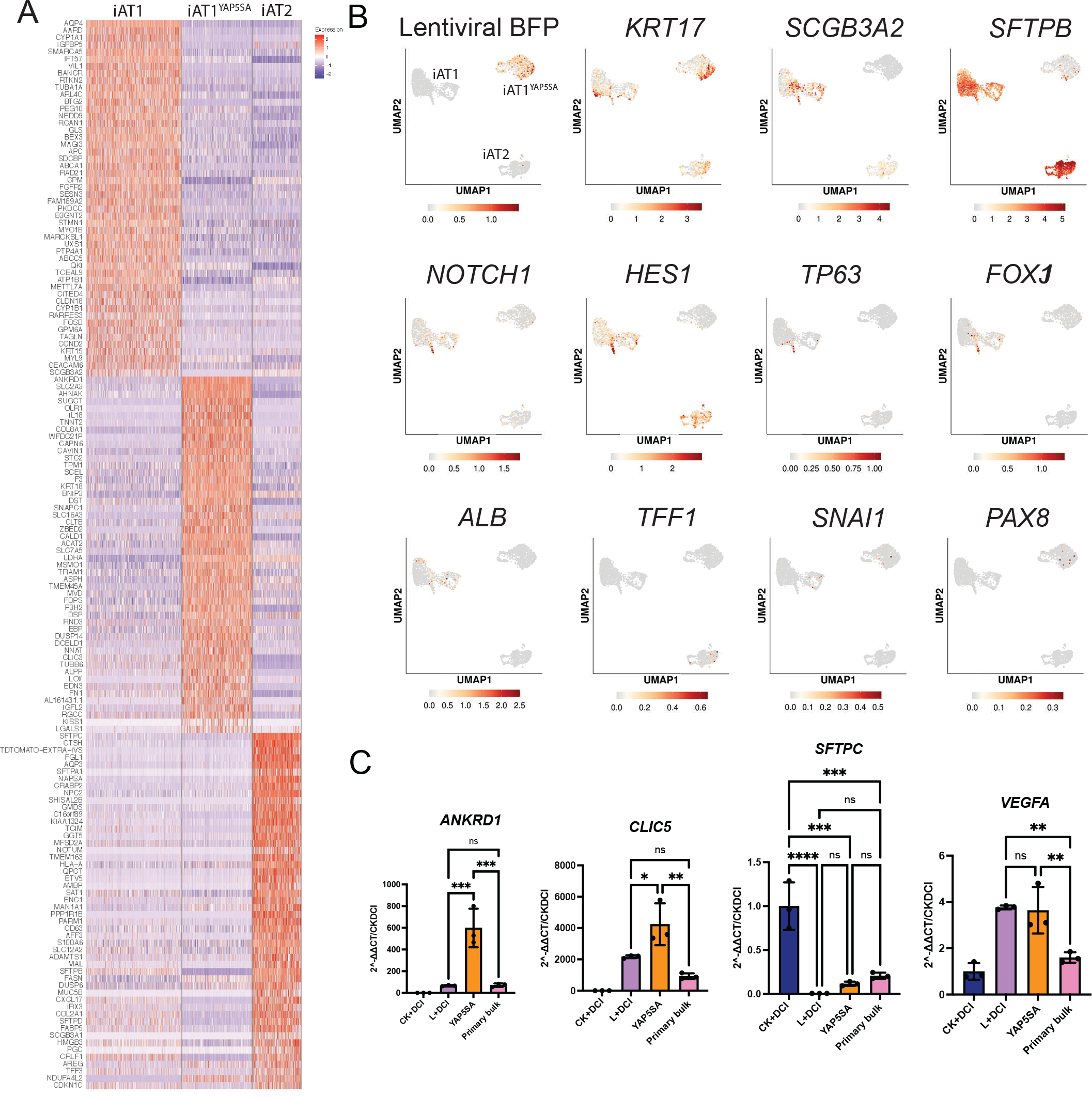
Single Cell RNA sequencing analysis of media and lentiviral induction of iAT1s. A) Heatmap showing top 50 differentially upregulated genes between iAT2, iAT1^YAP5SA^, and iAT1 populations. B) Gene expression overlays of lentiviral tagBFP, as well as transitional state markers, airway markers, and non-lung endoderm markers. C) Comparison of expression levels of indicated transcripts in whole well RNA extracts from iAT2, iAT1^YAP5SA^, and iAT1 populations compared to bulk primary human distal lung (RT-qPCR 2^-DDCt fold change compared to iAT2s in CK+DCI; N=3, 1-way ANOVA). p<0.05, **p<0.01, ***p<0.001, and ****p<0.001 for all panels.

**Supplemental Figure 9:**
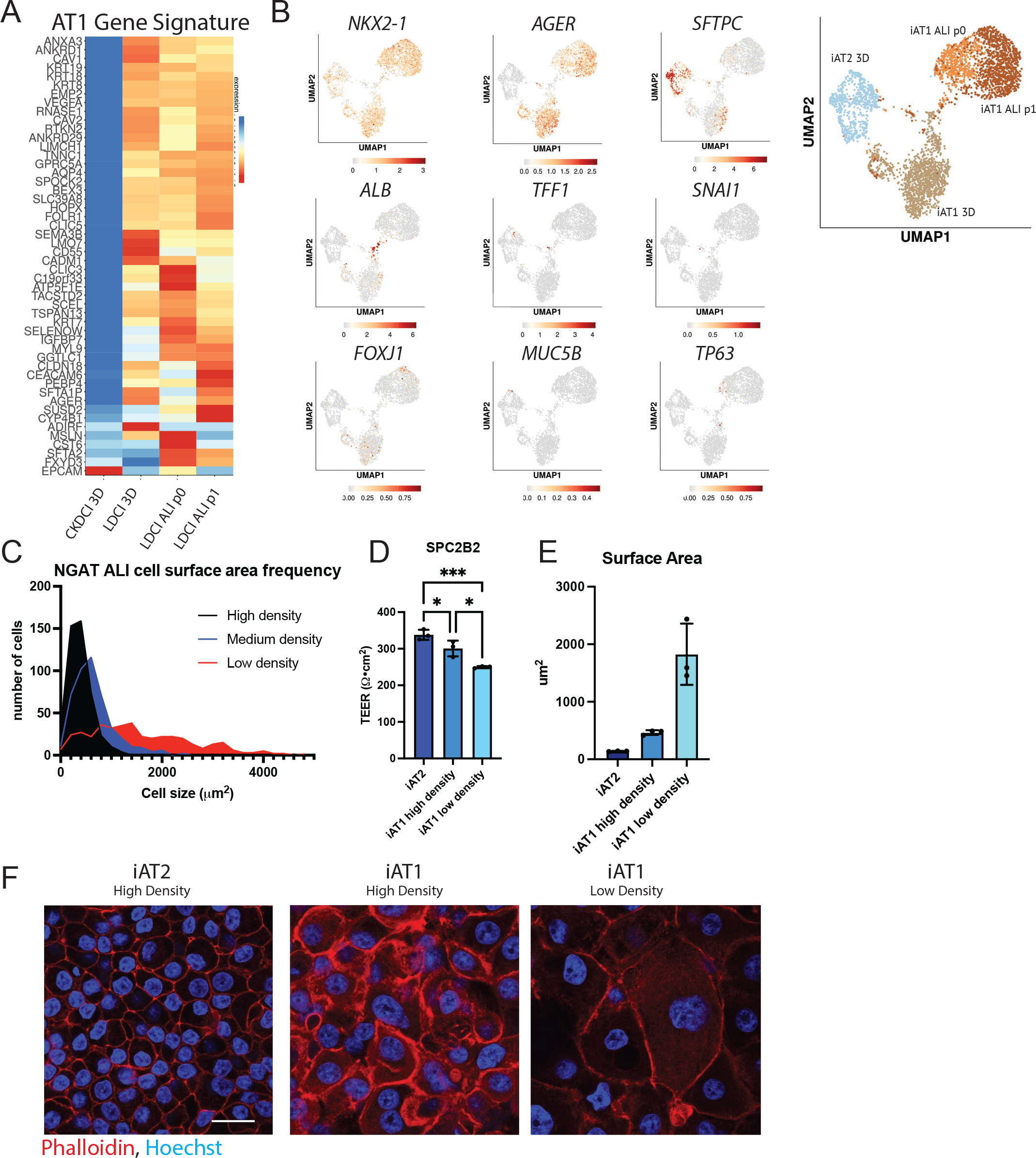
Profiling of iAT1s after Air Liquid Interface (ALI) Culture. A) Heatmap showing average expression of genes in 50 gene primary adult human AT1 gene signature across populations iAT2 3D, iAT1 3D, and iAT1 ALI conditions. B) UMAP expression overlays of lung epithelial markers and Non-lung markers. C) Frequency distribution of cell surface areas from high, medium, and low plating densities of iAT1s after ALI culture. Calculated using ImageJ on ZO-1 staining from ALIs fixed at day 10 post plating. N=457, 455, and 399 respectively across 3 different transwells. D) TEER of SPC2-ST-B2 iAT2s plated in CK+DCI after ALI culturing as published^68^ compared to iAT1s at high (200k) and low (50k) density plating in L+DCI and cultured at ALI. E) Surface area iAT2 vs iAT1 SPC2-ST-B2 N=3, averaged from ∼30- 50 cells per image. F) Immunofluorescence images of SPC2B2 iAT2s and iAT1s at high and low density. (F-actin: Phalloidin Red, Hoechst: blue) Scale bar = 20um. *p<0.05, **p<0.01, ***p<0.001, and ****p<0.001 for all panels.

